# Developmental exposure to a PFAS mixture impairs the anamnestic response to influenza A virus infection in mice

**DOI:** 10.1101/2025.11.10.687657

**Authors:** Christina M. Post, Jackie J. Agyemang, Katya A. McDonald, Carrie A. McDonough, B. Paige Lawrence

**Author notes:** CAM: Carnegie Mellon University, 4400 Fifth Avenue, Pittsburg, PA 15213, United States.

## Abstract

Developmental exposure to per- and polyfluoroalkyl substances (PFAS) has been linked to reduced antibody responses to childhood vaccines, but the underlying mechanisms remain unclear. Antibody production relies on interactions between various immune cell types, and it is unknown which are affected by PFAS exposure during development. To investigate this in a human-health relevant system, an in vivo model was established to delineate effects of developmental exposure to a mixture of four PFAS commonly found in human serum: PFOA, PFOS, PFHxS, and PFNA. Pregnant mice consumed water containing these PFAS throughout gestation and lactation. PFAS were measured in both mothers and offspring, and an exposure that avoided overt health issues was selected. The immune response to influenza A virus (IAV) infection was assessed in male and female offspring. Results showed that developmental PFAS exposure reduced IAV-specific antibody levels in both sexes. However, it diminished T follicular helper cells and germinal center B cells—critical for antibody production—in only female offspring. These findings highlight possible sex-specific immune effects and identify potential cellular mechanisms behind reduced antibody levels. Since these immune cells are essential for antibody production in humans, this study provides valuable insights into how PFAS exposure may impact human health.

**Synopsis:** A novel mouse model of developmental exposure to a human-relevant PFAS mixture recapitulates observations in epidemiological studies and also provides new insight into potential mechanisms of the lower antibody levels observed in humans.

## Introduction

Per- and polyfluoroalkyl substances (PFAS) are a large class of anthropogenic organic compounds that consist of either a fully or partially fluorinated carbon backbone attached to a functional group 1. PFAS have a wide range of properties that vary based on carbon chain length and functional group, which include being water and oil repellant and heat resistant, rendering them useful for numerous processes and products. PFAS are united in their ability to resist total environmental degradation and metabolism by living systems and because of this, they are often referred to as “forever chemicals” 2, 3. Widespread usage of PFAS in consumer products, packaging, mechanical, medical and electronic equipment, as well as firefighting foams, means that humans are regularly exposed to PFAS 4, 5. Recent reports from the National Health and Nutrition Examination Survey (NHANES) indicate that there are detectable levels of PFAS in the serum of ∼98% of the US population, including children, and half-lives in humans are estimated to range from 2-8 years 6-8.

Given the ubiquity of PFAS and their long half-lives, understanding the effects of exposure to PFAS has become a public health concern 9. Adverse health effects associated with PFAS exposure include thyroid disease, increased cholesterol, and liver and kidney cancer 5, 8, 10. PFAS also cross the placenta and are commonly detected in cord blood and breast milk, suggesting both in utero and lactational exposure occur 11-13. Environmental exposures that occur during early life development are particularly concerning because they have the potential to disrupt developmental processes and contribute to adverse health effects that persist throughout the lifespan 14-16. In fact, multiple epidemiological studies have demonstrated an association between developmental exposure to PFAS and reduced antibody titers to childhood vaccination 17-22.

In addition to being important for protection against infectious disease at both the individual and public health level, responses to vaccination provide a sentinel of a properly functioning immune system because antibody production involves many different immune cell types 23, 24. Vaccines are based on the principal of immunological memory, which enables the immune system to mount a rapid defense against a previously encountered antigen (i.e., vaccination or infection). Immunological memory is important for host defense against recurrent infections with commonly circulating pathogens such as influenza A viruses (IAV) 25. Whether elicited by natural infection or vaccination, the coordinated process that leads to the creation of pathogen-specific antibodies is referred to as humoral immunity.

How developmental PFAS exposure alters key cellular events that underpin the humoral immune response is not known. Establishing this is an essential first step to eventually decipher the mechanisms by which early life exposure to PFAS cause durable differences in immune system function. As such, a better understanding of how developmental exposure to PFAS influences immune responses has been highlighted as a critical need 26-28. Along with studies supporting connections between exposure to PFAS and immunotoxicity, it is becoming clear that most people are often exposed to more than one PFAS. Yet, most experimental studies of PFAS immunotoxicity examined exposure to a single PFAS, largely PFOS or PFOA 29-33. Thus, another identified need is studies that examine consequences of exposure to mixtures of PFAS 34. Among the myriad PFAS, PFOA, PFOS, PFHxS and PFNA are commonly co-detected in human cohorts in a range of geographic locations 8, 21, 26, 27, 35-37.

In the present study, we determined whether maternal exposure to a curated mixture of four PFAS influences the adaptive immune system in offspring using a mouse model and infection with human IAV. Since insufficient adaptive immunity can manifest as alterations in the response to primary infection and/or to repeated infection, we measured responses to primary infection and homotypic reinfection. Our findings are consistent with reports from epidemiological studies and provide new insight into how developmental exposure to PFAS perturbs humoral immune responses and immunological memory.

## Methods

### Developmental exposure

Adult (8-week-old) nulliparous female C57Bl/6 (B6) mice were paired with male B6 mice and were examined every morning for the presence of a vaginal plug. Adult B6 mice were purchased from The Jackson Laboratory (Bar Harbor, ME) and allowed to acclimatize for 1 week after arrival prior to mating. Female and male mice were separated and singly housed beginning on gestational day (GD) 0, which is the day of vaginal plug discovery. Impregnated female mice were randomly assigned to receive either plain vivarium drinking water (control, CTRL) or drinking water that had been spiked with the PFAS mixture (PFAS). Dams received either control or PFAS mixture-treated drinking water *ad libitum* throughout the entire period of gestation and lactation. That is, from GD0 until the offspring were weaned on postnatal day (PND) 21 (Figure 1A). This exposure strategy encompasses critical windows of immune system development, which begins *in utero* and continues postnatally ^38^. Cages were clearly labeled to denote treatment group, to ensure accurate dosing across pregnancy and postpartum. All members of the research team were aware of which animals were in which water treatment group. Water consumption and body weight of dams were monitored throughout the treatment period. Water bottles for all mice were changed weekly. At weaning (PND21), offspring were weighed, ear punched, and placed into cages with same sex littermates. No litters were culled. Single female offspring were co-housed with female offspring from the same developmental exposure group whenever possible. At weaning, all offspring received unspiked drinking water. Offspring were weighed weekly from weaning until they were infected with influenza A virus (IAV).

**Figure 1.**
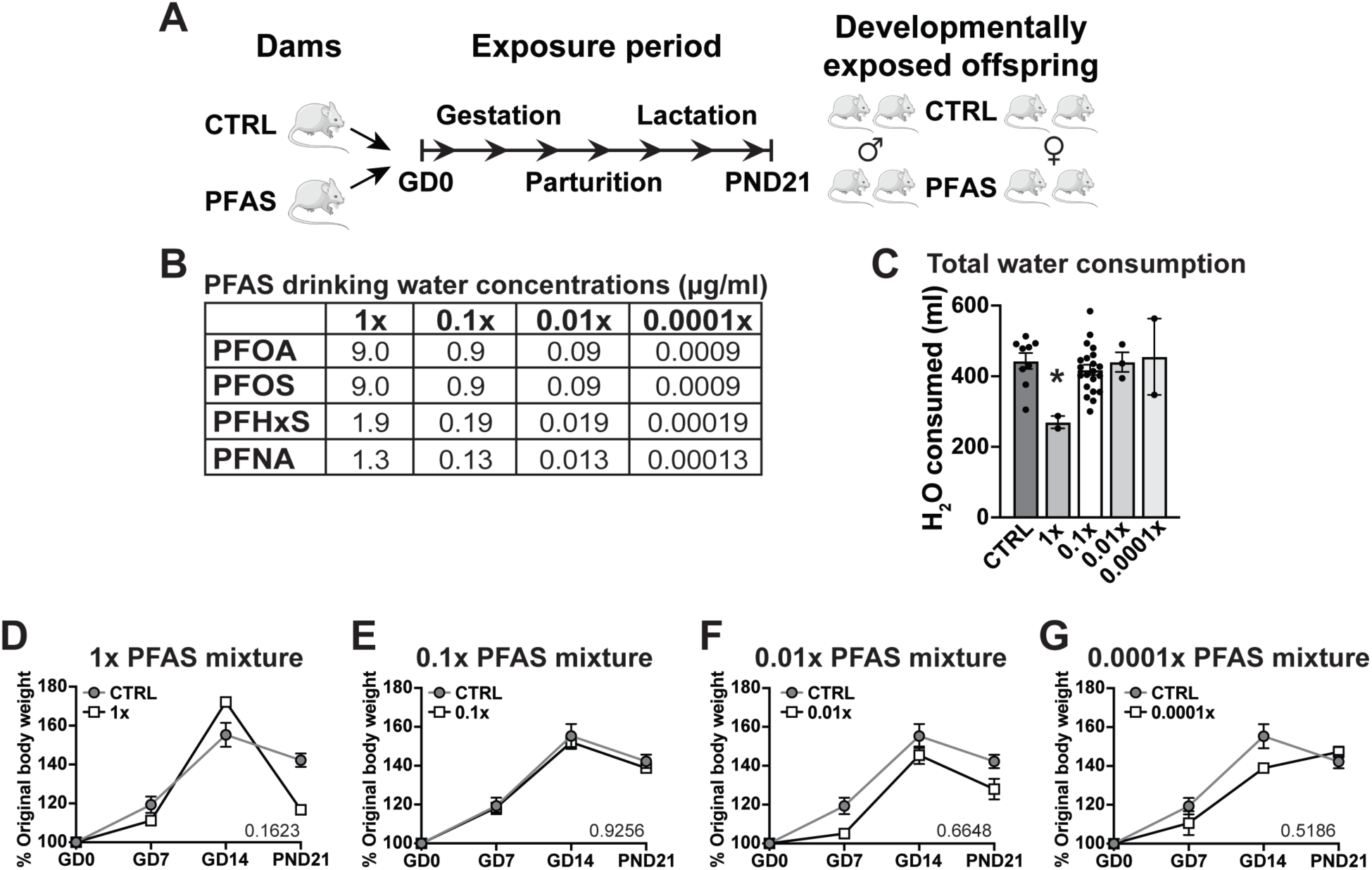
Developmental exposure strategy. (A) Dams were provided unspiked drinking water (control, CTRL) or drinking water spiked with a mixture of PFOA, PFOS, PFHxS, and PFNA (PFAS) continuously from gestational day (GD) 0 until the offspring were weaned on postnatal day (PND) 21. (B) Drinking water concentrations (µg/ml) of PFOA, PFOS, PFHxS, and PFNA in the 1x, 0.1x, 0.01x, and 0.0001x PFAS mixtures. (C) Mean (± SEM) total amount of water (ml) consumed by dams in each treatment group over the 40-day exposure period (gestation + lactation). Means were compared using a one-way ANOVA followed by Tukey’s HSD post hoc test. Asterisk (*) denotes p≤0.05 compared to control. (D-G) Dams were weighed weekly throughout gestation and on PND21, when the pups were weaned. Dams were not weighed between parturition and weaning to reduce stress. Mean body weight of dams that received the (D) 1x PFAS mixture, (E) 0.1x PFAS mixture, (F) 0.01x PFAS mixture, or (G) 0.0001x PFAS mixture compared to control dams. Data are presented as mean ± SEM and were analyzed by two-way ANOVA. The p-value for the interaction between treatment and day is provided on each graph. Data are from the following number of dams per group: Control n=9; 1x PFAS n=2; 0.01x PFAS n=21; 0.0001x PFAS n=2.

All mice were housed in microisolator cages in a specific-pathogen free facility at the University of Rochester, with a 12-hour light/dark cycle, ambient temperature between 20 and 22°C, and were provided food (LabDiet 5010) and water *ad libitum*. All animal treatments were conducted with prior approval of the Institutional Animal Care and Use Committee of the University of Rochester. The University has accreditation through the Association for Assessment and Accreditation of Laboratory Animal Care (AAALAC). Animals were treated humanely and with due consideration to the alleviation of any distress and discomfort. All guidelines from the U.S. Public Health Service Policy on Human Care and Use of Laboratory Animals were followed in handling of vertebrate animals.

### Preparation of PFAS mixture-spiked drinking water

Perfluorooctanoic acid (PFOA, CAS No. 335-67-1), perfluorooctane sulfonate (PFOS, CAS No. 2795-39-3), perfluorohexane sulfonate (PFHxS, CAS No. 3871-99-6), perfluorononanoic acid (PFNA, CAS No. 375-95-1) were each prepared as separate 100x stock solutions by dissolving the appropriate mass of the compound in deionized water. The concentration of each 100x PFAS stock solution was as follows: 1x PFAS mixture: 900 µg PFOA/ml, 900 µg PFOS/ml, 190 µg PFHxS/ml, 130 µg PFNA/ml; 0.1x PFAS mixture: 90 µg PFOA/ml, 90 µg PFOS/ml, 19 µg PFHxS/ml, 13 µg PFNA/ml; 0.01x PFAS mixture: 9 µg PFOA/ml, 9 µg PFOS/ml, 1.9 µg PFHxS/ml, 1.3 µg PFNA/ml; 0.0001x PFAS mixture: 0.09 µg PFOA/ml, 0.09 µg PFOS/ml, 0.019 µg PFHxS/ml, 0.013 µg PFNA/ml. Concentrated stock solutions were handled using aseptic technique and stored in glass bottles at 4°C. Fresh dosing solutions were prepared weekly as a master mix by combining each stock solution with vivarium drinking water (sourced from the Monroe County, NY Water Authority) then aliquoted into polycarbonate mouse drinking water bottles. All PFAS were purchased from Sigma-Aldrich (St. Louis, MO) and were ≥95% pure. PFOS and PFHxS were purchased as potassium salts and concentrations were calculated using the total compound mass. Based on levels commonly detected in human tissues and blood, the curated quaternary mixture contained equal concentrations of PFOA and PFOS, 5-times less PFHxS and 7-times less PFNA ^6, 7, 39^. The use of drinking water as a route of exposure is relevant to how humans are exposed to PFAS ^5^. The final concentrations of PFAS in the drinking water were as follows: 0.0009 to 9 µg PFOA/ml, 0.0009 to 9 µg PFOS/ml, 0.00019 to 1.9 µg PFHxS/ml, and 0.00013 to 1.3 µg PFNA/ml. Based on mice with an average body weight (BW) of 20 g that consume an average of 4 ml H_2_O/day, these concentrations in the water administer an estimated daily dose of 0.18 to 1880 µg PFOA/kg BW, 0.18 to 1880 µg PFOS/kg BW, 0.0376 to 376 µg PFHxS/kg BW, and 0.026 to 260 µg PFNA/kg BW. Mice drink more water and gain weight as pregnancy progresses, and we acknowledge the uncertainty this adds to the estimated daily dose.

### Influenza A virus infection

Adult (≥ 8-week-old) offspring were anesthetized via intraperitoneal (i.p.) injection with 225-250 µL avertin (2% 2,2,2-tribromomethanol; Sigma-Aldrich) then infected intranasally (i.n.) with 120 hemagglutination units (HAU) influenza A virus (IAV) strain A/HKx31 (HKx31; H3N2) diluted in 25 µl endotoxin-tested sterile phosphate-buffered saline (PBS, pH 7.4). This inoculum is sublethal in immunocompetent mice ^40–42^. To examine whether developmental exposure to the PFAS mixture affected the anamnestic response to IAV, some mice were reinfected with 120 HAU of HKx31 72 days after primary infection. The initial stock of influenza A/HKx31 was provided by Dr. M. Coppola (Argonex, Charlottesville, VA), and was propagated titered and stored at -80°C as previously described ^41, 43^. In all experiments, infection and tissue collection were performed in the morning. All work with infectious agents was conducted with prior approval of the Institutional Biosafety Committee of the University of Rochester, following guidelines of the US NIH/CDC.

### Tissue collection

Prior to tissue collection, mice were euthanized by i.p. injection with a lethal dose of Euthasol (Virbac, Greeley, CO) followed by exsanguination. Blood was collected by cardiac puncture and serum was separated by centrifugation at 8000 x g for 10 minutes at 4°C and then stored at -80°C. Spleen and liver were harvested at weaning, weighed then flash frozen in liquid nitrogen prior to storage at -80°C. Mediastinal lymph nodes (MLN) were removed on days 0, 9, 14, and 70 relative to primary infection and on days 3 and 5 relative to reinfection. On each day, single cell suspensions of MLN cells from individual mice were prepared by using the frosted ends of microscope slides to disrupt the tissue and release the cells. The cells were resuspended in cold Hanks Balanced Salt Solution (HBSS) containing 2.5% fetal bovine serum (HBSS/FBS; Hyclone, Logan, UT). Red blood cells were lysed at room temperature by 5 min incubation with a buffer consisting of 0.15 M NH_4_Cl, 10 mM NaHCO_3_, and 1 mM EDTA. The addition of cold HBSS/FBS was used to quench the lysis, and the cell suspension was passed through a 70 µm filter to remove debris. The cell suspensions were centrifuged (200 x g, 10 min, 4°C) to pellet cells, which were resuspended in cold HBSS/FBS. The number of cells and cellular viability were determined using trypan blue dye and a TC20^TM^ cell counter (BioRad, Hercules, CA).

### Analytical flow cytometry

Single cell suspensions of freshly prepared MLN cells from individual animals were stained for flow cytometric analysis. Antibody information, including vendor, catalog number, clone, and amount used is listed in Table S1. The following three separate staining panels were used: (1) B cells, (2) Tfh cells and CD8+ T cells, and (3) CD4+ T helper subsets (Th1, Th2, Th17, Treg). For each staining panel, 2 x 10^6^ cells from each mouse were incubated with anti-mouse CD16/32 antibodies (clone 93) for 10 min to block nonspecific staining. An antibody cocktail targeting extracellular antigens was added directly to each sample. Following incubation for 20 min at 4°C, cells were washed twice with staining buffer (PBS, 1% BSA, 0.01% sodium azide). Washing consisted of centrifugation (200 x g, 2 min, 4°C), removal of supernatant and resuspension in buffer. Samples stained with the B cell panel were immediately fixed with PBS containing 2% formaldehyde. For samples in the Tfh cell and CD8+ T cell panel, cells were incubated with PE conjugated streptavidin for 30 minutes and washed again prior to fixation with 2% formaldehyde. For enumeration of CD4+ T helper subsets, extracellular staining was followed by 30-minute fixation/permeabilization using the FoxP3 staining kit (Invitrogen, Carlsbad, CA). Nonspecific intracellular staining was blocked by incubating cells again with anti-mouse CD16/32. Cells were incubated a cocktail of antibodies against lineage-specific transcription factors for 20 min, washed twice with the FoxP3 staining kit buffer then fixed with 2% formaldehyde. All staining was performed at 4°C, and during incubation periods cells were maintained in the dark. Prior to use, antibodies were titrated to determine the optimal concentrations. Fluorescence minus one (FMO) controls were used to define gating parameters and determine nonspecific fluorescence. To exclude debris and dead cells, gating on singlets and light scatter properties [forward scatter (FSC) and side scatter (SSC)] were performed prior to gating with cell-specific markers. Samples were run on a BD LSRII flow cytometer (BD Biosciences, San Jose, CA) and data was analyzed using FlowJo v10 (TreeStar, Ashland, OR).

### Anti-influenza virus antibody ELISA

Relative levels of influenza virus-specific antibodies in serum were measured by enzyme-linked immunosorbent assay (ELISA), as previously described ^44–46^. Briefly, flat bottom ELISA plates (Greiner Bio-One MICROLON™) were coated with 5 µg/ml purified IAV strain HKx31 (Charles River, Wilmington, MA) overnight at 37°C. Nonspecific binding was blocked by incubating plates with 5% BSA in PBS for 2 h at room temperature. The plates were washed with PBS containing 0.05% Tween 20 in between each step. Plates were incubated with serially diluted serum (1:100 to 1:819,200 in 5% BSA/PBS) overnight at 4°C. Biotinylated goat anti-mouse IgM and IgG antibodies (Southern Biotechnology, Birmingham, AL) were diluted to 1:5000 in 5% BSA/PBS and plates were incubated at room temperature for 45 min. Avidin-peroxidase (1:400 dilution in 5% BSA/PBS, Sigma-Aldrich, St. Louis, MO) was added and plates were incubated for 30 min at room temperature. The colorimetric substrate [2,2’-azino-bis(3-ethylbenz-thiazo-line-6-sulfonic acid) with 0.03% H_2_O_2_] was added and the plates were developed at room temperature. A SpectraMax Plate reader (Molecular Devices, San Jose, CA) was used to read absorbance values at 405 nm. To measure the avidity of IAV-specific IgG, plates were incubated with 1.0 M guanidine hydrochloride (GuHCl; Sigma-Aldrich, St. Louis, MO) in buffer consisting of PBS, 0.2% Tween 20, and 10 mg/ml BSA for 15 min at room temperature prior to incubation with biotinylated IgG ^45, 47, 48^.

### Serum PFAS measurements

Mouse serum was thawed to room temperature and vortexed thoroughly. A 100 µl aliquot of each serum sample was added to an Agilent Enhanced Matrix Removal (EMR) Lipid cartridge (Agilent Technologies, Santa Clara, CA) along with 25 µl of a mass-labeled PFAS mixture (50 ng/ml) in MeOH and 400 µl of crash solvent (0.1 M formic acid in cold acetonitrile). The mixture was allowed to passively mix for ten minutes then eluted under ultrahigh purity nitrogen on a positive pressure manifold. The eluent was collected in polypropylene vials and 25 µl of injection standard (^13^C-labeled PFOA in MeOH, 50 ng/ml) was added. Samples were prepared alongside triplicate solvent blanks (100 µl water) and matrix blanks (100 µl bovine calf serum). All extracts were refrigerated until analysis.

Extracts were injected (35 µl volume) onto an Agilent liquid chromatography system coupled to an Agilent 6560 quadrupole-time-of-flight mass spectrometer (LC-QTOF-MS) using a similar analytical method as described in detail in a previous publication ^46^. Concentrations of PFOA, PFOS, PFHxS, and PFNA were measured based on response factors from calibration curves analyzed in the same run. PFOS was measured as the summed linear and branched isomers. All calibration curves were required to include ≥ 5 calibration points and to have a linear fit with r^2^ ≥ 0.99. The calibration range was defined as the concentration range for which measured concentration was within ±30% of known concentrations. The method reporting limit for each analyte was established as the minimum point included in the calibration unless the analyte was found in the blank at a greater level, in which case it was the average method blank plus three standard deviations. Continuing calibration verification (CCV) samples with known concentrations of target analytes spiked into bovine serum were analyzed after every ten samples and were within ±30% of expected concentrations in all cases. All analytical standards, including mass-labeled internal standards, injection standards, and native standards, were obtained from Wellington Laboratories (Guelph, Ontario, Canada). All solvents and reagents were LC-MS-grade (Fisher Scientific, Hampton, NH).

### Real-time quantitative PCR

Total RNA was extracted from liver tissue using a Qiagen RNeasy RNA isolation kit with on-column DNAse (Qiagen, Germantown, MD) digestion of genomic DNA. Complementary DNA (cDNA) was synthesized from 1 µg total RNA using the BioRad iScript™ cDNA synthesis kit (Hercules, CA). qPCR reactions were assembled using the BioRad iQ™ SYBR^®^ Green Supermix and run on a BioRad CFX96 Touch Real-Time PCR Detection System (Hercules, CA). Gene expression of acyl-CoA oxidase 1 (*Acox1,* Accession No. AF006688) relative to the housekeeping gene *Gapdh* was calculated using the 1′1′Ct method ^49^. The forward (FW) and reverse (RV) primer sequences for *Acox1* were as follows: *Acox1* FW: 5’-GGGAGTGCTACGGGTTACATG-3’ and RV: 5’-CCGATATCCCCAACAGTGATG-3’ ^50^.

### Statistical analysis

In these experiments, the dams received treatment; hence, the dams and not the offspring are the statistical unit. With the exception of the data presented in Figure 2, which includes data from all pups, the group size denotes the number of dams. For the IAV infection time course, at each point in time relative to infection, same sex offspring were from different dams. Except for area under the curve (AUC) analysis, which was performed with Prism software (v.10.4.1, GraphPad Software, Boston, MA), all data were analyzed using JMP (v.17.2.0, SAS Institute Inc., Cary, NC). Differences between two groups across multiple points in time were compared using a two-way analysis of variance (ANOVA) followed by either a Tukey’s Honest Significance Difference (HSD) or Dunnett’s post-hoc test. Differences between four groups at a single point in time were compared by one-way ANOVA followed by Tukey’s HSD post-hoc test. Differences in means were considered statistically significant at p ≤ 0.05. Error bars on all graphs represent SEM. The specific statistical test used is indicated in each figure legend.

**Figure 2.**
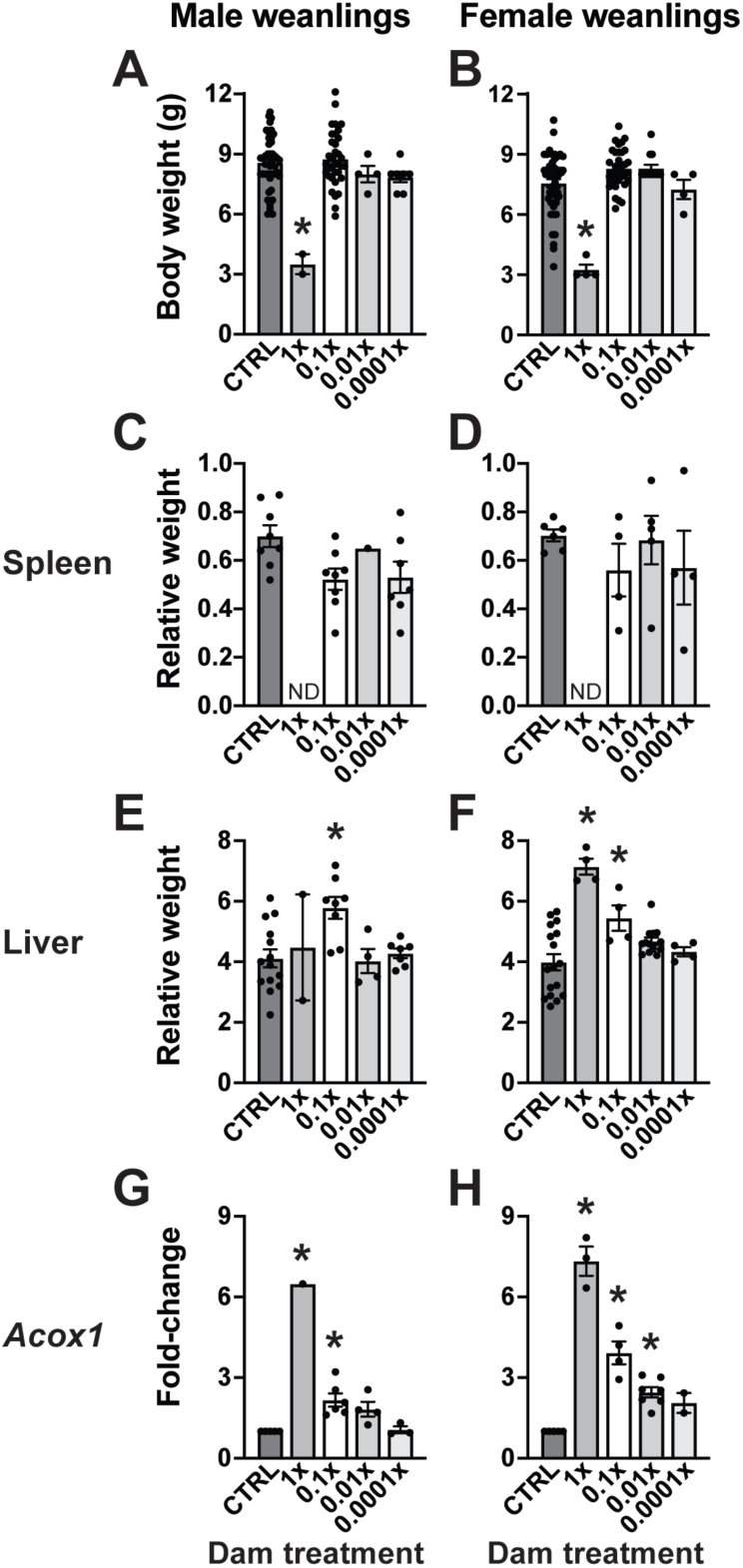
Offspring body weight, relative spleen and liver weight, and Acox1 expression at weaning. All offspring were weighed on the day that they were weaned. Data are from all viable weanlings in each exposure group. (A) Mean body weight of male offspring (n=2-47). (B) Mean body weight of female offspring (n=4-49). For some experiments, spleen and liver were removed at weaning. Organ weights are expressed as a ratio of body weight. (C) Mean spleen weight of male offspring (n=1-8). (D) Mean spleen weight of female offspring (n=4-6). (E) Mean liver weight of male offspring (n=2-14). (F) Mean liver weight of female offspring (n=4-17). qPCR was used to measure Acox1 expression in the livers of (G) male offspring (n=1-6) and (H) female offspring (n=2-7). Error bars on all graphs represent mean ± SEM. Data were analyzed by one-way ANOVA followed by Tukey’s HSD post hoc test. Asterisk (*) denotes p≤0.05 compared to control within each sex. Numerical values and a complete list of p-values are provided in Table S3. ND: not determined.

## Results

### Dose selection for developmental immunotoxicity assessment

To ascertain whether developmental exposure to PFAS affected the adaptive immune system later in life, we exposed pregnant mice via the drinking water to a mixture of four PFAS that are commonly detected in human tissues (PFOA, PFOS, PFHxS and PFNA) ^6, 7, 27, 35, 51, 52^ (Figure 1A). Control mice received unspiked drinking water. Prior to undertaking immunological assessments in the offspring, initial studies were conducted to identify a concentration of this quaternary PFAS mixture that would not adversely impact pregnancy success or cause overt toxicity in the pups. This was necessary because neither a no observable adverse effect level (NOAEL) nor a lowest observable adverse effect level (LOAEL) for PFAS developmental immunotoxicity has been established for individual PFAS or mixtures of PFAS. However, a NOAEL for suppressed IgM production following direct exposure to PFOA during adulthood has been reported ^53^. This provided a benchmark for the highest dose exposure group, which was given drinking water containing 9 µg PFOA/ml, 9 µg PFOS/ml, 1.9 µg PFHxS/ml, and 1.3 µg PFNA/ml. PFAS concentrations in this curated mixture were not equivalent because PFAS levels in human serum and tissues are generally not equivalent. The concentrations of PFOA and PFOS in human serum tend to be similar while in several reports, the concentration of PFHxS was roughly 5-fold lower than that of PFOA/PFOS and the concentration of PFNA was about 7 times lower than PFOA/PFOS ^6, 7, 27, 39^. The proportion of PFOA, PFOS, PFHxS and PFNA relative to each other was maintained and separate groups of dams were exposed to 10-fold lower (0.1x), 100-fold lower (0.01x), or 10,000-fold lower (0.0001x) concentrations of PFAS (Figure 1B).

PFAS concentrations in serum collected from dams showed a dose-related pattern (Table 1). Dams that received the 1x PFAS mixture consumed significantly less water during the treatment period than dams that received control water; however, water consumption by dams treated with the other (lower concentration) PFAS mixtures was not significantly different than that of control dams (Figure 1C). In all PFAS exposure groups, dam body weight increase over the course of pregnancy did not differ significantly different from control dams (Figure 1D-G). There were also no differences in time to parturition, litter size, or sex ratio between offspring of control and PFAS-treated dams (Table S2).

**Table 1.** Serum PFAS concentrations (ng/ml) in dams.

|  | **Control water**<br>**(n=4)** |  | **1x PFAS water<sup>a</sup>**<br>**(n=2)** |  | **0.1x PFAS water<sup>b</sup>**<br>**(n=4)** |  | **0.01x PFAS water<sup>c</sup>**<br>**(n=3)** |  | **0.0001x PFAS water<sup>d</sup>**<br>**(n=2)** |  |
| --- | --- | --- | --- | --- | --- | --- | --- | --- | --- | --- |
|  | **Mean** | **SD** | **Mean** | **SD** | **Mean** | **SD** | **Mean** | **SD** | **Mean** | **SD** |
| **PFOA** | 0.88 | 0.24 | 38247<sup>f</sup> | 4355 | 26383<sup>f</sup> | 5276 | 1090 | 241 | 38.33 | 2.66 |
| **PFOS<sup>e</sup>** | 15.07 | 7.40 | 87745<sup>f</sup> | 13439 | 34368<sup>f</sup> | 10483 | 1265 | 289 | 78.87 | 21.26 |
| **PFHxS** | 0.82 | 0.32 | 21125<sup>f</sup> | 3048 | 23048<sup>ef</sup> | 6494 | 695 | 118 | 21.94 | 5.90 |
| **PFNA** | 1.13 | 0.28 | 12010 | 1815 | 6547<sup>g</sup> | 1760 | 182 | 35 | 9.26 | 1.09 |

Serum was collected from dams on the day pups were weaned (i.e., PND21). PFAS concentrations were measured by LC-HRMS. SD: standard deviation.
<sup>a</sup>1x PFAS mixture: 9 µg/ml PFOA, 9 µg/ml PFOS, 1.9/ml µg PFHxS, 1.3 µg/ml PFNA
<sup>b</sup>0.1x PFAS mixture: 0.9 µg/ml PFOA, 0.9 µg/ml PFOS, 0.19 µg/ml PFHxS, 0.13 µg/ml PFNA
<sup>c</sup>0.01x PFAS mixture: 0.09 µg/ml PFOA, 0.09 µg/ml PFOS, 0.019 µg/ml PFHxS, 0.013 µg/ml PFNA
<sup>d</sup>0.0001x PFAS mixture: 0.0009 µg/ml PFOA, 0.0009 µg/ml PFOS, 0.00019 µg/ml PFHxS, 0.00013 µg/ml PFNA
<sup>e</sup>Sum of linear and branched isomers.
<sup>f</sup>All values greater than highest calibration curve point.
<sup>g</sup>Two of four values greater than highest calibration curve point.

While pregnancy outcomes were not adversely affected by exposure to the highest concentration of the PFAS mixture, the mean body weight of male and female offspring at weaning was significantly reduced compared to same-sex offspring of control dams. Male offspring of dams treated with the 1x PFAS mixture weighed 59% less than control male offspring (Figure 2A), and female weanlings weighed 57% less than female offspring of control dams (Figure 2B). In contrast, the body weight of weanlings in the lower PFAS concentration groups was not significantly different from same-sex control offspring. At weaning, the relative spleen weight of male (Figure 2C) and female (Figure 2D) offspring was not affected by developmental exposure to the three lower concentrations of the PFAS mixture. The relative liver weight of male offspring of dams that drank the 0.1x PFAS mixture was increased 1.4-fold compared to male offspring of control dams (Figure 2E). Compared to female control offspring, there was a 1.8-fold increase in the relative liver weight of female offspring of dams that received the 1x PFAS mixture and a 1.4-fold increase in female offspring of dams that consumed the 0.1x PFAS mixture (Figure 2F).

At the time of weaning, offspring of dams exposed to the 0.1x, 0.01x, and 0.0001x PFAS mixtures had PFOA, PFOS, PFHxS and PFNA levels that were greater than offspring of dams that received control water (Table 2). The concentration of each PFAS was progressively less with declining concentration in the dam’s drinking water, indicating a dose-related pattern in offspring serum PFAS concentration. Due to their small size, we were unable to obtain a sufficient volume of serum from offspring of dams in the 1x PFAS mixture water group. However, in prior studies, direct exposure to PFOA and PFOS induced hepatic expression of the fatty acid oxidation gene acyl-CoA oxidase 1 (Acox1) 46, 54-56. Therefore, livers were harvested at weaning to assess this biochemical metric of PFAS exposure in all groups of offspring. In male weanlings, compared to control, developmental exposure to the 1x PFAS mixture caused a 6.5-fold increase in Acox1 expression, developmental exposure to the 0.1x PFAS mixture caused a 2-fold increase, while the 0.01x and 0.0001x PFAS mixtures did not induce Acox1 expression (Figure 2G). In female weanlings, maternal consumption of the 1x, 0.1x, and 0.01x PFAS mixtures elicited statistically significant and dose-dependent elevation of Acox1 expression compared to control (Figure 2H). Based on these collective assessments, dams in all subsequent studies were given drinking water spiked with the 0.1x PFAS mixture.

**Table 2.** Serum PFAS concentrations (ng/ml) in developmentally exposed offspring at weaning.

| **Male offspring** |  |  |  |  |  |  |  |  |
| --- | --- | --- | --- | --- | --- | --- | --- | --- |
|  | **Control water**<br>(n=4) |  | **0.1x PFAS water<sup>a</sup>**<br>(n=4) |  | **0.01x PFAS water<sup>a</sup>**<br>(n=3) |  | **0.0001x PFAS water<sup>a</sup>**<br>(n=2) |  |
|  | **Mean** | **SD** | **Mean** | **SD** | **Mean** | **SD** | **Mean** | **SD** |
| **PFOA** | 1.23 | 1.00 | 4212<sup>c</sup> | 1335 | 416 | 128 | 16.33 | 2.69 |
| **PFOS<sup>b</sup>** | 5.93 | 1.25 | 7246 | 2727 | 345 | 56 | 23.14 | 6.36 |
| **PFHxS** | 1.17 | 0.85 | 4493<sup>b</sup> | 599 | 237 | 51 | 11.01 | 1.98 |
| **PFNA** | 0.78 | 0.14 | 1471 | 264 | 90 | 22 | 4.29 | 0.87 |
| **Female offspring** |  |  |  |  |  |  |  |  |
|  | **Control water**<br>(n=4) |  | **0.1x PFAS water<sup>a</sup>**<br>(n=3) |  | **0.01x PFAS water<sup>a</sup>**<br>(n=3) |  | **0.0001x PFAS water<sup>a</sup>**<br>(n=2) |  |
|  | **Mean** | **SD** | **Mean** | **SD** | **Mean** | **SD** | **Mean** | **SD** |
| **PFOA** | 1.12 | 0.79 | 3540<sup>c</sup> | 609 | 404 | 116 | 17.36 | 1.76 |
| **PFOS<sup>b</sup>** | 7.07 | 1.22 | 5482 | 853 | 348 | 104 | 26.00 | 0.74 |
| **PFHxS** | 1.95 | 1.36 | 3523<sup>b</sup> | 238 | 231 | 60 | 10.64 | 1.35 |
| **PFNA** | 1.06 | 0.22 | 1024 | 68 | 86 | 23 | 5.06 | 0.81 |

Serum was collected from developmentally exposed offspring at weaning (PND21) and PFAS concentrations were measured by LC-HRMS. SD: standard deviation.
<sup>a</sup>Denotes relative concentration of PFAS mixture in water consumed by the dam. All offspring received unspiked (control) water at weaning.
<sup>b</sup>Sum of linear and branched isomers.
<sup>c</sup>Three of four values higher than highest calibration curve point.

#### Assessment of T cell responses to IAV following developmental PFAS exposure

The effects of developmental exposure to a mixture of PFOA, PFOS, PFHxS, and PFNA on the primary and memory response to IAV infection were examined in adult offspring (Figure 3A). Developmental exposure to this PFAS mixture did not affect the body weight of male or female offspring prior to infection or after acute primary IAV infection compared to same-sex offspring of control dams (Figure S1). Mediastinal lymph nodes (MLN), which drain the lower respiratory tract, were removed to directly interrogate cellular aspects of the adaptive immune response to IAV. In response to IAV infection, the number of MLN cells increases, contracts after recovery, and then, upon re-infection increases again 57, 58. These infection-induced changes in cellularity did not differ between same-sex offspring of control and PFAS-treated dams at any point in time after primary or recall infection (Table S4).

**Figure 3.**
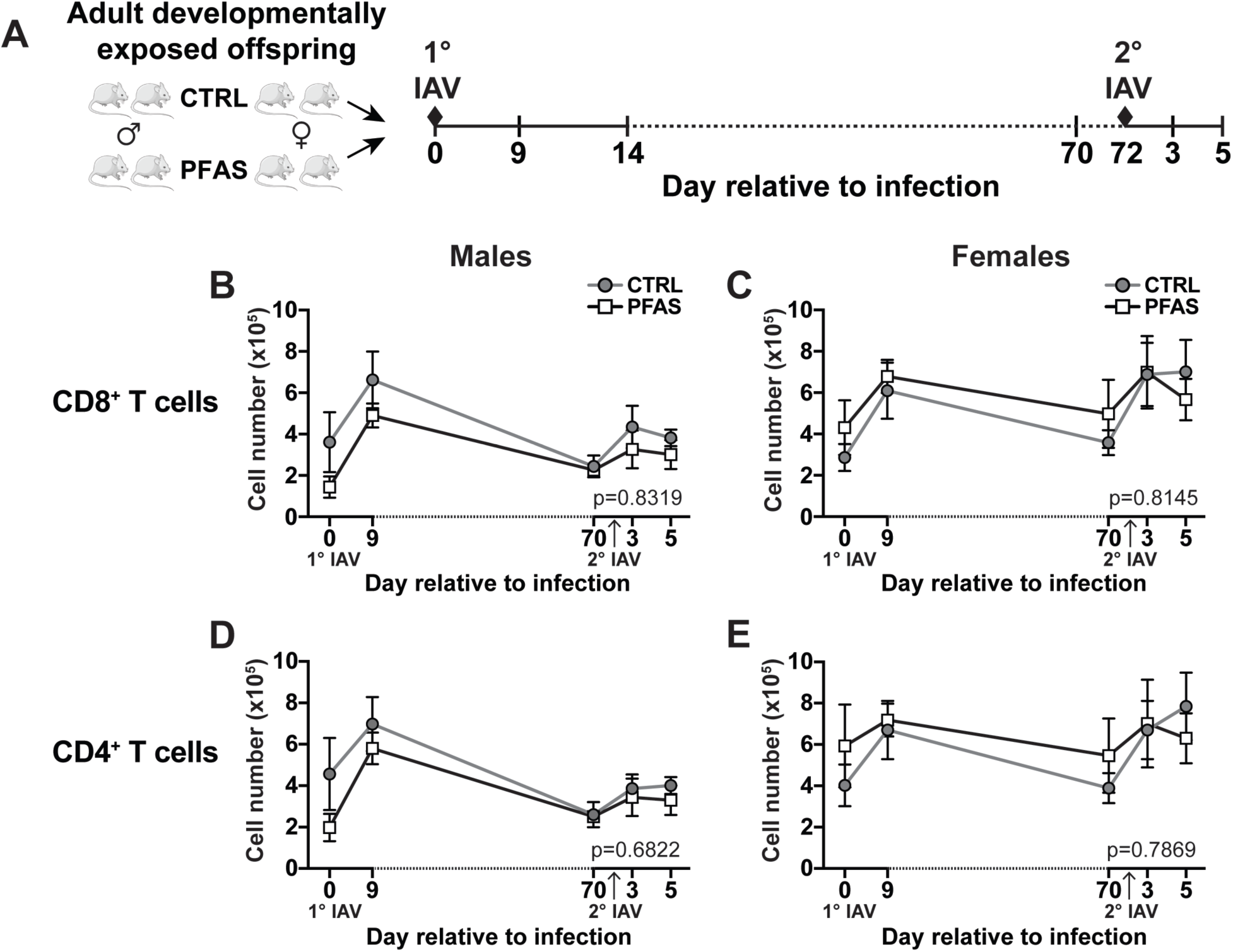
Developmental exposure to the PFAS mixture did not perturb the number of CD8+ or CD4+ T cells. (A) Adult (8-week-old) male and female offspring of control and PFAS-treated dams were infected with influenza A virus (IAV, strain HKx31). Mediastinal lymph nodes (MLNs) were harvested on days 9, 14, and 70 after primary infection, and 3 and 5 days after re-infection. The re-infection with HKx31 was performed 72 days after primary infection. A separate group of mice were not infected, and data from these mice are denoted as “Day 0” relative to infection. Graphs depict the mean number of CD8+ T cells in the MLN of (B) male offspring and (C) female offspring and the mean number of CD4+ T cells in the MLN of (D) male offspring and (E) female offspring during primary and secondary IAV infection. On each day relative to infection there were 4-8 male and 3-6 female mice per group. At each point in time, same sex offspring were from different dams. Data are represented as mean ± SEM and were analyzed by two-way ANOVA followed by Tukey’s HSD post hoc test. The p-value for the interaction between treatment and day relative to infection is provided on each graph. Numerical values and a complete list of p-values are provided in Table S4.

In response to infection, the number of CD8+ and CD4+ T cells similarly increases, contracts and increases in response to re-infection 25, 57, 58. In both males (Figure 3B, Table S4) and females (Figure 3C, Table S4), there was no difference in the number of CD8+ T cells in offspring of control and PFAS-treated dams at any point in time examined. Developmental exposure to this PFAS mixture also did not significantly modulate the total number of CD4+ T cells in the MLN of male (Figure 3D, Table S4) or female (Figure 3E, Table S4) offspring on any day relative to infection compared to same-sex control offspring.

Given that CD4+ T cells are not homogeneous, we further examined whether early life PFAS exposure affected the distribution of functionally distinct subsets prior to or after infection. Th1 and T follicular helper (Tfh) cells are the main CD4+ T cell subsets involved in the response to IAV 59-61. Th1 cells support viral clearance through secretion of antiviral cytokines that attract and activation of other immune cells 60. While the percentage and number of Th1 cells was altered by infection, developmental exposure to the PFAS mixture did not further affect the percentage (Table S6) or number (Table S7) of Th1 cells in male or female offspring at any point in time relative to infection.

Tfh cells provide essential signals to B cells in germinal centers. These interactions support plasma cell differentiation, antibody affinity maturation, antibody isotype switching, and regulate immunological memory 62, 63. In male offspring of control dams, the percentage of Tfh cells increased in response to primary infection and was significantly higher on days 9 and 70 compared to day 0 (Figure 4A, Table S6). However, in male offspring of PFAS-treated dams, the percentage of Tfh cells was not significantly different from uninfected mice on either day 9 or day 70 (Figure 4A, Table S6). The number of Tfh cells in male offspring of control and PFAS-treated dams exhibited similar changes in response to infection, peaking on day 9, then returning to baseline (Figure 4B, Table S7). There was also a significant increase in the percentage of Tfh cells from female control offspring on day 9 post-primary infection compared to day 0; however, in female PFAS offspring, the percentage of Tfh cells was blunted, and not significantly different on day 9 compared to uninfected controls (Figure 4C, Table S6). The number of Tfh cells in female control offspring increased significantly after primary infection on day 9, returned to baseline by day 70, then increased in response to secondary infection on day 5 (Figure 4D, Table S7). Conversely, in female offspring of PFAS-treated dams, the number of Tfh cells from infected mice was not significantly greater than that of uninfected mice on any day relative to infection (Figure 4D, Table S7).

**Figure 4.**
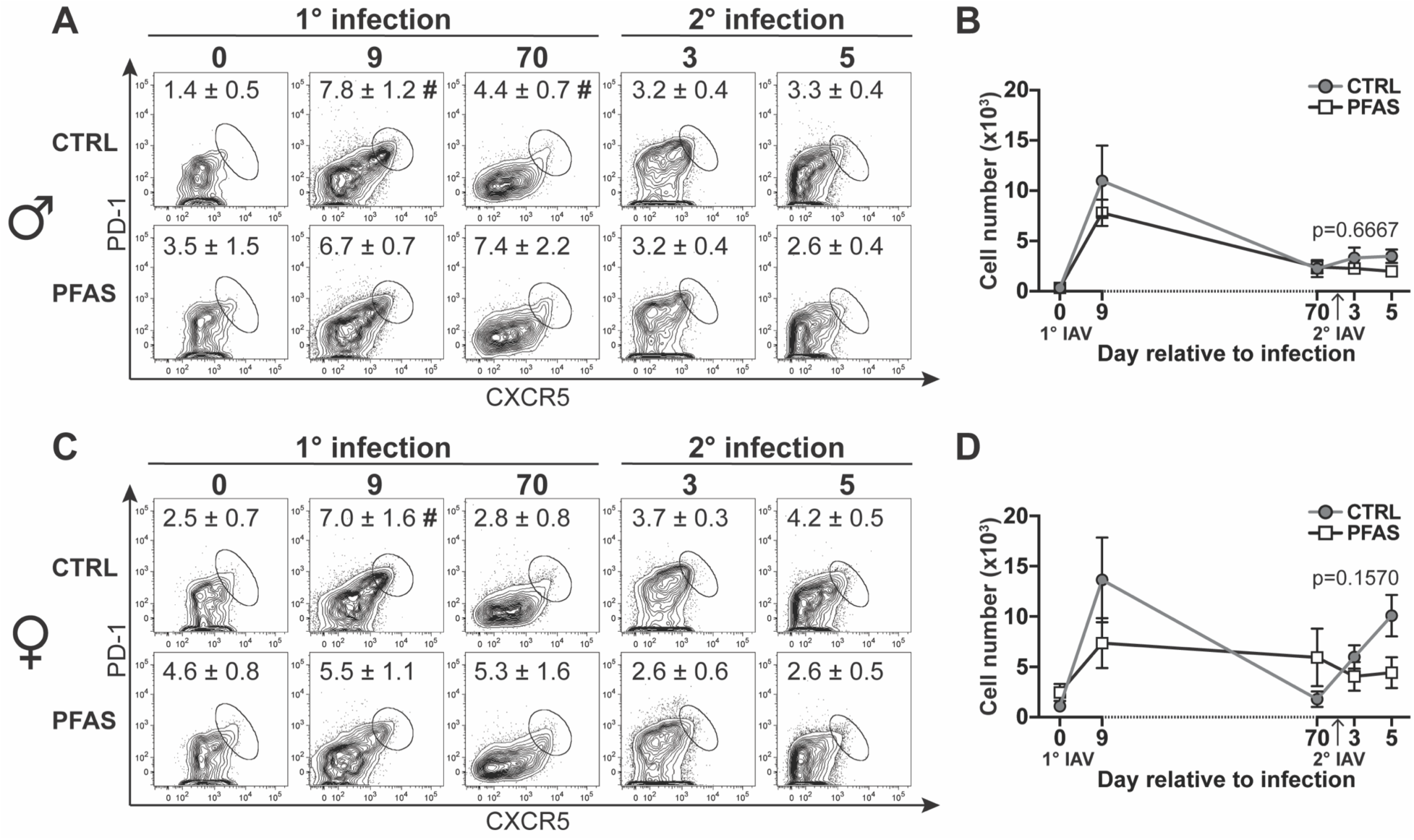
Developmental exposure to the PFAS mixture alters the Tfh cell response to IAV. Offspring of control and PFAS-treated dams were infected with IAV, and MLNs were harvested as depicted in Figure 3A. Flow cytometry was used to analyze Tfh cells (CD4+CD44hiCXCR5+PD-1hi) after primary and secondary IAV infection. The gating strategy used to identify Tfh cells is shown in Figure S2. (A) Representative FACS plots depict the percentage of Tfh cells in male offspring on the indicated days relative to infection. The number on each plot is the mean percentage of CD44hiCD4+ T cells identified as Tfh cells. (B) The mean number of Tfh cells in male offspring on the indicated day relative to infection. (C) Representative FACS plots depict the percentage of Tfh cells in the MLN of female offspring. The number on each plot is the mean percentage of CD44hiCD4+ T cells that are Tfh cells. (D) The mean number of Tfh cells in female offspring on the indicated day relative to infection. On each day relative to infection, there were 4-8 male and 3-6 female offspring per group. At each point in time, same-sex offspring were from different dams. Data are represented as mean ± SEM and were analyzed by two-way ANOVA followed by Dunnett’s and Tukey HSD post hoc tests. # symbol denotes p≤0.05 calculated by Dunnett’s test comparing day relative to infection to day 0 within sex/dam water treatment group. The p-value for the interaction between treatment and day relative to infection is provided on each graph. Numerical values and complete lists of p-values are provided in Table S6 (percentage of Tfh cells) and Table S7 (number of Tfh cells).

In addition to Th1 and Tfh cells, the percentage and number of the Th2, Th17, and T regulatory (Treg) cell CD4+ T cell subsets were enumerated prior to and after IAV infection. With one exception, developmental exposure to this PFAS mixture did not affect the percentage (Table S6) or number (Table S7) of Th2, Th17, or Treg cells in the MLN of either sex on any day relative to infection compared to same-sex control offspring. The exception was that the percentage of Th17 cells in uninfected female PFAS offspring was four times higher than uninfected offspring of control dams. However, this difference did not persist after infection. Thus, in female offspring, developmental exposure to the PFAS mixture had the most notable impact on the Tfh cell response to both primary and secondary IAV infection.

#### Developmental exposure to PFAS mixture dampened aspects of the B cell response

Germinal center (GC) B cells play a central role in humoral immunity. In response to signals from Tfh cells, they undergo class switch recombination and somatic hypermutation to create antibodies with higher affinity and can develop into plasma cells and memory B cells 64. In male offspring of control dams, compared to immunologically naïve mice, the percentage of GC B cells in the MLN was significantly higher 14 and 70 days after primary IAV, and 5 days after reinfection (Figure 5A, Table S8). In male offspring of PFAS-treated dams, the overall pattern was similar. The percentage of GC B cells was greater on day 14 post-primary infection and day 5 post-reinfection than it was in uninfected mice (day 0). However, unlike control offspring, the percentage of GC B cells 70 days after primary infection was not significantly different from uninfected mice (Figure 5A, Table S8). In male offspring, the number of GC B cells showed a similar pattern of change in response to first and second infection, but there was no statistically significant difference between the exposure groups (Figure 5B, Table S8).

**Figure 5.**
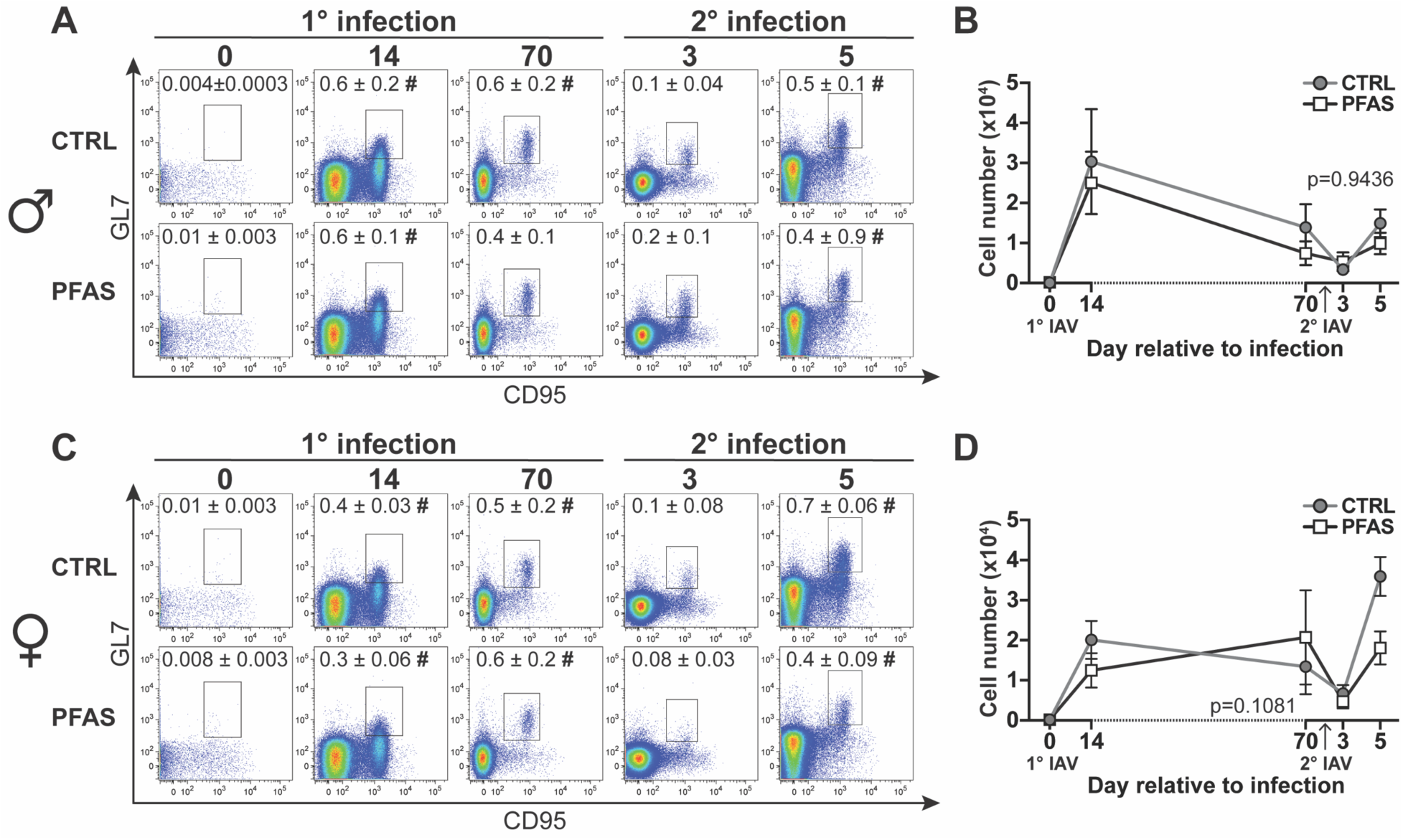
The effects of developmental exposure to the PFAS mixture on germinal center (GC) B cells in the MLN. Offspring of control and PFAS-treated dams were infected with IAV, and MLNs were harvested as depicted in Figure 3A. Flow cytometry was used to analyze GC B cells (CD3- B220+CD95+GL7+) in the MLN after primary and secondary IAV infection. The gating strategy used to identify GC B cells is shown in Figure S3. (A) Representative FACS plots depict the percentage of GC B cells in male offspring on the indicated days relative to infection. The number on each plot is the mean percentage of MLN cells that were identified as GC B cells. (B) The mean number of GC B cells from male offspring on the indicated day relative to infection. (C) Representative FACS plots depict the percentage of GC B cells in female offspring. The number on each plot is the mean percentage of MLN cells that were GC B cells. (D) The mean number of GC B cells in female offspring on the indicated day relative to infection. On each day relative to infection, there were 4-7 male offspring and 3-6 female offspring per group. At each point in time, same-sex offspring are from different dams. Data are depicted as mean ± SEM and were analyzed by two-way ANOVA followed by Dunnett’s and Tukey HSD post hoc tests. # symbol denotes p≤0.05 calculated by Dunnett’s test comparing day relative to infection to day 0 within sex/dam water treatment group. The p-value for the interaction between treatment and day relative to infection is provided on each graph. Numerical values and complete lists of p-values are provided in Table S8.

In female offspring from both developmental exposure groups, the percentage of GC B cells in the MLN was significantly increased on day 14 and day 70 after primary infection and on day 5 post-secondary infection compared to day 0 (Figure 5C, Table S8). The number of GC B cells from female offspring of control dams increased on day 14 post-primary infection, returned to baseline by day 70, then increased again in response to secondary infection on day 5 (Figure 5D, Table S8). Unlike control, the number of GC B cells from female PFAS offspring on day 14 post-primary infection was not greater than that of uninfected mice, however there were more GC B cells in the infected mice on day 70 (Figure 5D, Table S8). Similar to control, GC B cells from female PFAS offspring also increased 5 days after reinfection. The effect on GC B cells was not due to differences in the total B cell population as there were no differences in the number (Table S4) or percentage (Table S5) of B cells in the MLN between same-sex offspring of control and PFAS-treated dams at any time point relative to infection. We also compared the percentage and number of plasma cells in MLNs from control and PFAS offspring during primary and secondary IAV infection and found that there were no significant differences in plasma cells between control and PFAS offspring on any day relative to infection in either sex (Table S8).

#### IAV-specific antibody levels

Infection with IAV elicits the production of virus-specific antibodies. Fourteen days after primary infection, there were no discernable differences in the levels of circulating IAV-specific antibodies in male (Figure 6A) or female (Figure 6B) offspring from control and PFAS exposed dams. However, 5 days after reinfection with the same virus, male offspring of PFAS-treated dams had 27% less IAV-specific IgG than male offspring of control dams (Figure 6A, Table S9). In female offspring of PFAS-treated dams, the relative level of IAV-specific IgG was 16% lower than that of female control offspring 70 days after primary infection and 21% lower 5 days after reinfection (Figure 6B, Table S9). Developmental exposure to the PFAS mixture did not influence IAV-specific IgM levels in either male (Figure S4A, Table S9) or female (Figure S4B, Table S9) offspring on any day relative to infection compared to same-sex offspring of control dams.

**Figure 6.**
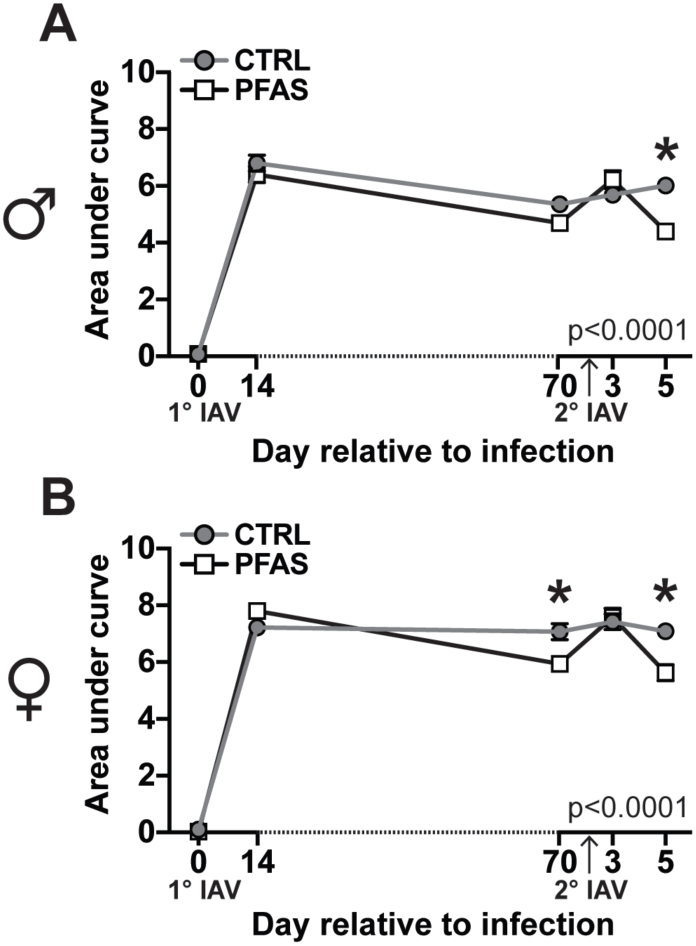
IAV-specific IgG levels during primary and secondary infection. Male and female offspring of control and PFAS-treated dams were infected with IAV as depicted in Figure 3A. Serum was collected at the indicated time points relative to infection and relative levels of circulating IAV-specific IgG were measured by ELISA and area under curve (AUC) analysis was performed. Graphs depict mean AUC values of IAV-specific IgG levels throughout primary and secondary IAV infection in (A) male offspring (n=3-6 mice per time point) and (B) female offspring (n=4-6 mice per time point). Data are represented as mean ± SEM and were compared by two-way ANOVA followed by Tukey’s HSD post hoc test. The p-value for the interaction between treatment and day relative to infection is provided on each graph. A full list of AUC values and p-values is provided in Table S9. Asterisk (*) denotes p≤0.05 for PFAS vs. CTRL on the indicated day relative to infection.

To assess whether developmental exposure to this PFAS mixture affected the overall quality of virus-specific antibodies, we used a modified ELISA to determine relative antibody avidity 45, 47. In male offspring of control treated dams, treatment with 1M GuHCl caused a 23% reduction in virus-specific IgG, and this decrease was observed across the dilution series compared to that of untreated serum (Figure 7A). In male offspring of PFAS-treated dams, treatment with GuHCl reduced IAV-specific IgG levels 25% compared to untreated serum (Figure 7B). Therefore, despite having lower levels of IAV-specific IgG than control offspring, the relative avidity of IgG from male PFAS offspring for IAV was similar to that of control offspring (Figure 7C). Treatment with 1M GuHCl did not significantly reduce IAV-specific IgG levels in serum from female offspring of control dams (Figure 7D, 7F). Conversely, when serum from female PFAS offspring was treated with GuHCl, there was a significant 16% reduction in IAV-specific IgG levels compared to untreated serum (Figure 7E, 7F). These results indicate that in addition to affecting the amount of virus-specific IgG in both male and female offspring, developmental exposure to this PFAS mixture reduced the relative strength with which IAV-specific IgG from female offspring binds to antigen, i.e., the avidity of the antibody.

**Figure 7.**
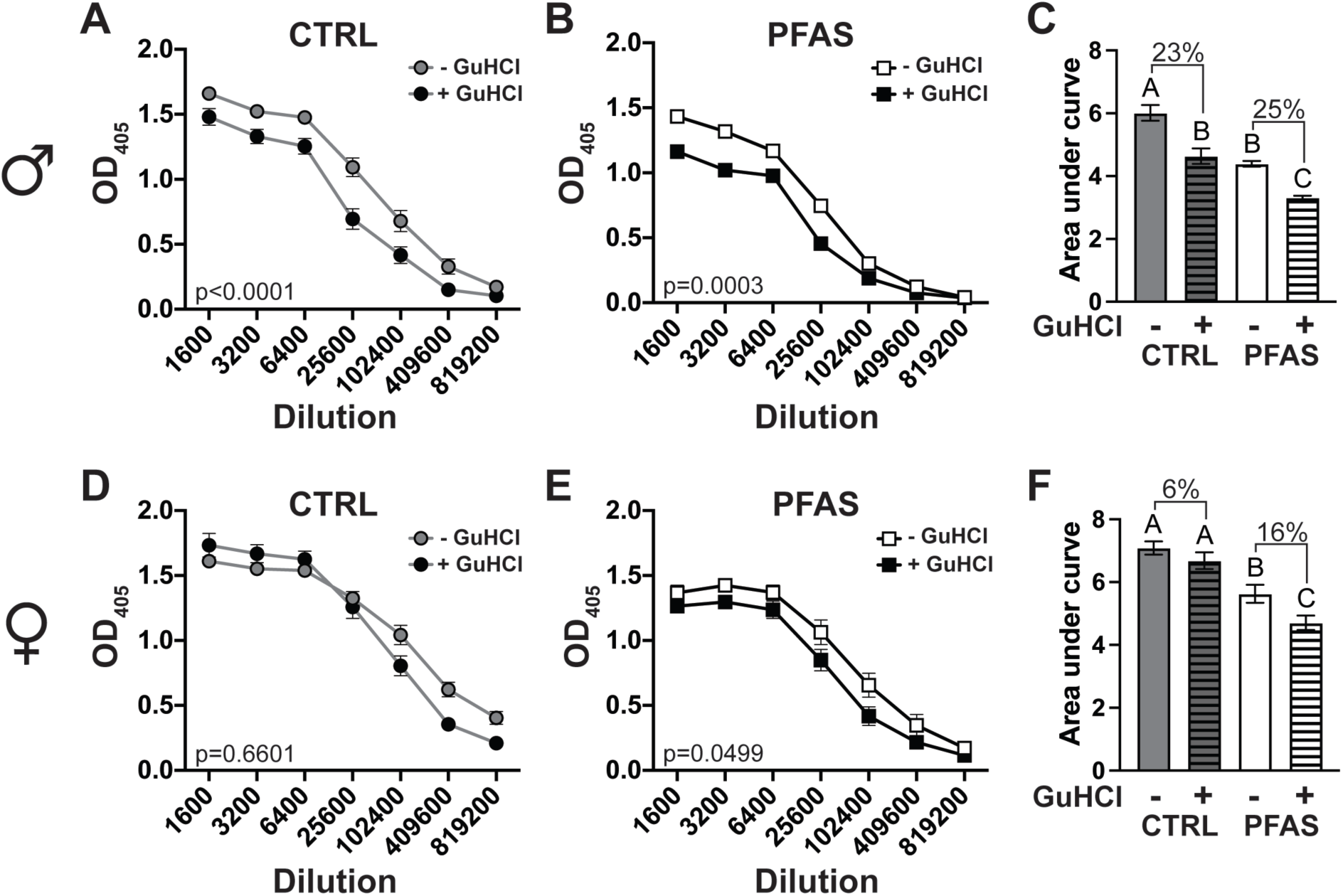
Avidity of IAV-specific IgG on day 5 post-reinfection. Serum was collected from male and female offspring 5 days after reinfection with IAV and the relative avidity of IAV-specific IgG was assessed by ELISA. (A, B) IAV-specific IgG levels in serially diluted serum from male offspring of (A) control dams (n=6) and (B) PFAS-treated dams (n=6) that was either untreated or treated with 1M GuHCl. (C) Bar graph depicts the mean AUC for the line graphs in panels A and B. The percentages denote the reduction in AUC caused by treatment with 1M GuHCl. (D, E) IAV-specific IgG levels in serially diluted serum from female offspring of (D) control dams (n=5) and (E) PFAS-treated dams (n=5) that was either untreated or treated with 1M GuHCl. (F) Bar graph depicts the mean AUC for line graphs in panels D and E. The percentages denote the reduction in AUC caused by treatment with 1M GuHCl. Data are represented as mean ± SEM. For panels C and F, means were compared by one-way ANOVA followed by Tukey’s HSD post hoc test. Bars that do not share the same letter designation are significantly different from one another (p≤0.05). p-values for all comparisons are listed in Table S10.

## Discussion

Recent assessments of adverse health effects associated with PFAS consistently emphasize that the immune system is particularly sensitive 26, 65, 66. For instance, population-based studies support that exposure to PFAS may contribute to immunosuppression, such as increased susceptibility to infection 67 and reduced responses to vaccines 19. Epidemiological studies also suggest that fetal or early life periods may be especially sensitive to PFAS, but few experimental studies have empirically defined the relationship between developmental exposure to PFAS and differences in immune responses 68. This is critically important because experimental and epidemiological studies have shown that exposures experienced during developmentally sensitive periods, such as in the womb or immediately after birth, even at levels that do not perturb the fully mature immune system, can impact immune responses later in life 14-16. In the presented study, we used a mixture of four PFAS that are commonly detected in human fluids and tissues and determined whether developmental exposure affected key metrics of the primary and memory immune response to infection with IAV. We report that developmental exposure to this PFAS mixture perturbed some, but not all, aspects of the adaptive immune response to IAV, particularly after the second infection, and that the consequences were different in male and female offspring. Our findings are broadly consistent with epidemiological studies, which have consistently reported associations between higher maternal serum PFAS levels and lower antibody titers to childhood vaccination 19, 21, 26, 37. Moreover, we have extended understanding of the immunological consequences of developmental PFAS exposure. Based on the endpoints measured, and the PFAS mixture and strain of virus used, the impact of developmental exposure to PFAS was more pronounced in female offspring and the humoral response appeared more affected than the cell-mediated response. These new findings further reveal that the impact of developmental exposure to PFAS extends beyond the relative level of circulating antibodies produced and may affect the quality of antibodies. Avidity refers to the strength of binding between antibodies and cognate antigens, and in female offspring of PFAS-treated dams the overall avidity of IAV-specific IgG for antigen was reduced. While antibody avidity has not been interrogated in human cohort studies, similar cellular mechanisms influence antibody affinity maturation in mice and humans 69. Therefore, this finding suggests that developmental exposure to PFAS may affect human immunity by diminishing both the quantity and quality of protective antibodies elicited in response to infection or vaccination.

Our findings also reveal that the mechanism by which PFAS affects humoral immunity is likely multifactorial, as developmental exposure to the PFAS mixture diminished the frequency of Tfh cells and germinal center B cells in female offspring upon reinfection; however, IgG levels were reduced in both sexes. Therefore, it is unlikely that differences in percent and number of Tfh cells and germinal center B cells fully explain the reduction in virus-specific IgG levels. Tfh cells interact with B cells in germinal centers in lymphoid organs. Germinal centers provide a critical niche for B cell affinity maturation, clonal selection, differentiation, and maturation of plasma cells, which are the main source of secreted antibodies, and memory B cells 62, 64. Developmental exposure to the PFAS mixture used in this study did not affect the percentage or number of plasma cells. Consequently, the decrease in virus-specific IgG levels in PFAS exposed offspring is not simply the result of fewer plasma cells. We also did not observe any differences in IAV-specific IgM levels between offspring of control and PFAS-treated dams, which suggests that developmental exposure to the PFAS mixture may interfere with B cell class switching, a process that requires Tfh cell help 62. Therefore, one cellular level mechanism by which developmental exposure to PFAS reduces antibody responses could be via modulation of the bidirectional communication between Tfh cells and B cells that supports robust humoral immunity. Defects in the signaling capacity or function of either Tfh cells, germinal center B cells, or both could impair plasma cell development and/or function and hinder antibody secretion. It is also possible that developmental exposure to PFAS affects the accessory cells that support T cells and B cells, such as dendritic cells, monocytes and macrophages.

Findings from the mouse model developed and used in this study are consistent with results from separate studies in which exposure to PFAS was associated with alterations in affected human Tfh cells and B cells 27, 70, 71. Specifically, the diminution in Tfh cells observed in developmentally exposed mice is similar to a recently reported inverse association between maternal PFAS levels and the number of Tfh cells in infants 27. In another recent report, children exposed to PFAS had lower levels of rotavirus-specific antibodies, which was associated with reduced Tfh cell cytokine secretion 71. Our findings are also consistent with a transcriptomic analysis of peripheral blood mononuclear cells from adults exposed to PFAS in which genes and pathways related to germinal center formation, germinal center reactions, and plasma cell development were downregulated by PFAS 70. Experimental studies using exposure of adult mice to either PFOA or PFOS have also reported decreased antibody production and decreases in T cell and B cell subsets 72-74, and we previously demonstrated that germinal center B cells were reduced in adult mice exposed directly to the 1x PFAS mixture 46. Decreased antibody production and alterations in B cells have also been reported in a study of developmental exposure to PFOS. In that study, IgM production was decreased in adult male, but not female, offspring while B cell frequency was reduced in female offspring only 75.

Sex differences in PFAS immunotoxicity have been consistently reported across experimental animal studies; however whether males or females are more affected varies, and may depend on the endpoint being examined 33, 72, 74-76. For the endpoints examined in this study, female offspring appeared more sensitive than male offspring to the effects of developmental exposure to the PFAS mixture. Male and female offspring of PFAS-treated dams had similar concentrations of each of the four PFAS in their serum, therefore differences in PFAS body burden do not provide a simple explanation of sex differences. In both humans and rodents, there are inherent biological sex differences in the immune response to influenza virus infection, with females typically mounting a greater immune response to infection and vaccination 58, 77, 78. Thus, the greater magnitude of the immune response to IAV in females could make the effects of PFAS exposure more pronounced. Many different factors contribute to sex differences in immune responses; however, one intriguing possibility that relates specifically to PFAS is the role of peroxisome proliferator-activated receptors (PPARs). PFAS activation of PPARs, specifically PPARα and/or PPARγ, has been proposed as a molecular initiating event in the PFAS adverse outcome pathway 54, 79. In humans and mice, PPARα and PPARγ expression on T cells and B cells begins during embryonic development 80-82. Expression of PPARα and PPARγ on T cells is sexually dimorphic, with PPARα being more highly expressed in T cells from males and PPARγ more highly expressed in T cells from females 83. Interestingly, activation of PPARγ has been shown to suppress Tfh cells and reduce germinal center formation specifically in female mice 84. Therefore, it is possible that sex differences in the effects of PFAS exposure could be due to differences in the activation of PPARs or other receptors in males and females.

Demonstrating that maternal exposure to a mixture of PFAS to which humans are commonly exposed alters adaptive immune response to a common human respiratory pathogen provides a deeper understanding of cellular events perturbed by these ubiquitous chemicals. While it is challenging to perfectly mimic human exposure in experimental model systems, this study demonstrates that developmental immunotoxicity occurred at internal doses in mice that are within the range of levels detected in human population-based studies 6, 7, 51. The cellular mechanisms responsible for humoral immunity are similar in rodents and humans. Given that developmental PFAS exposure of mice recapitulated findings from human cohort studies, this offers an experimental system to define mechanisms that is readily translatable to human health. Tfh cells and well-regulated antibody production are central to not only responses to infection and vaccination but are key mediators in autoimmune disease 85. Therefore, the health implications extend beyond the ability of PFAS to disrupt host defenses against pathogens. Early life exposure to PFAS may influence immune-mediated diseases more broadly.

## Supporting information

Offspring post-primary infection body weight data, representative gating strategies, flow cytometry antibodies, pregnancy outcomes, and tables of numerical values and p-values that correspond to the figures in the text (PDF)

## Acknowledgements

We would like to thank Dr. Timothy Bushnell and Matt Cochran at the Center for Advanced Research Technologies (CART) at the University of Rochester for the flow cytometry resources.

## Author contributions

CMP: designed and performed experiments, analyzed data, interpreted results, prepared figures, drafted manuscript. JJA: performed experiments. KAM: performed experiments. CAM: performed experiments, analyzed data. BPL: conceived project and designed experiments, interpreted results, edited and revised manuscript. CMP, JJA, KAM, CAM, and BPL approved manuscript for submission.

## References

(1) Buck, R. C.; Franklin, J.; Berger, U.; Conder, J. M.; Cousins, I. T.; de Voogt, P.; Jensen, A. A.; Kannan, K.; Mabury, S. A.; van Leeuwen, S. P. Perfluoroalkyl and polyfluoroalkyl substances in the environment: terminology, classification, and origins. Integr Environ Assess Manag 2011, 7 (4), 513–541. DOI: 10.1002/ieam.258 From NLM Medline.

(2) Wang, Z.; DeWitt, J. C.; Higgins, C. P.; Cousins, I. T. A Never-Ending Story of Per- and Polyfluoroalkyl Substances (PFASs)? Environ Sci Technol 2017, 51 (5), 2508–2518. DOI: 10.1021/acs.est.6b04806 From NLM Medline.

(3) Cousins, I. T.; DeWitt, J. C.; Gluge, J.; Goldenman, G.; Herzke, D.; Lohmann, R.; Ng, C. A.; Scheringer, M.; Wang, Z. The high persistence of PFAS is sufficient for their management as a chemical class. Environ Sci Process Impacts 2020, 22 (12), 2307–2312. DOI: 10.1039/d0em00355g From NLM Medline.

(4) De Silva, A. O.; Armitage, J. M.; Bruton, T. A.; Dassuncao, C.; Heiger-Bernays, W.; Hu, X. C.; Karrman, A.; Kelly, B.; Ng, C.; Robuck, A.;, et al. PFAS Exposure Pathways for Humans and Wildlife: A Synthesis of Current Knowledge and Key Gaps in Understanding. Environ Toxicol Chem 2021, 40 (3), 631–657. DOI: 10.1002/etc.4935 From NLM Medline.

(5) Sunderland, E. M.; Hu, X. C.; Dassuncao, C.; Tokranov, A. K.; Wagner, C. C.; Allen, J. G. A review of the pathways of human exposure to poly-and perfluoroalkyl substances (PFASs) and present understanding of health effects. J Expo Sci Environ Epidemiol 2019, 29 (2), 131–147. DOI: 10.1038/s41370-018-0094-1 From NLM Medline.

(6) Calafat, A. M.; Kato, K.; Hubbard, K.; Jia, T.; Botelho, J. C.; Wong, L. Y. Legacy and alternative per- and polyfluoroalkyl substances in the U.S. general population: Paired serum-urine data from the 2013-2014 National Health and Nutrition Examination Survey. Environ Int 2019, 131, 105048. DOI: 10.1016/j.envint.2019.105048 From NLM Medline.

(7) CDC. Fourth national report on human exposure to environmental chemicals: updated tables, January 2019, Volume one; Centers for Disease Control and Prevention, 2019. https://stacks.cdc.gov/view/cdc/75822/cdc_75822_DS1.pdfDOI: 10.15620/cdc75822.

(8) Fenton, S. E.; Ducatman, A.; Boobis, A.; DeWitt, J. C.; Lau, C.; Ng, C.; Smith, J. S.; Roberts, S. M. Per- and Polyfluoroalkyl Substance Toxicity and Human Health Review: Current State of Knowledge and Strategies for Informing Future Research. Environ Toxicol Chem 2021, 40 (3), 606–630. DOI: 10.1002/etc.4890 From NLM Medline.

(9) Rosato, I.; Bonato, T.; Fletcher, T.; Batzella, E.; Canova, C. Estimation of per- and polyfluoroalkyl substances (PFAS) half-lives in human studies: a systematic review and meta-analysis. Environ Res 2024, 242, 117743. DOI: 10.1016/j.envres.2023.117743 From NLM Medline.

(10) ATSDR. *Toxicological profile for Perfluoroalkyls*; Agency for Toxic Substances and Disease Registry, Atlanta, GA, 2021. DOI: 10.15620/cdc:59198.

(11) Mamsen, L. S.; Bjorvang, R. D.; Mucs, D.; Vinnars, M. T.; Papadogiannakis, N.; Lindh, C. H.; Andersen, C. Y.; Damdimopoulou, P. Concentrations of perfluoroalkyl substances (PFASs) in human embryonic and fetal organs from first, second, and third trimester pregnancies. Environ Int 2019, 124, 482–492. DOI: 10.1016/j.envint.2019.01.010 From NLM Medline.

(12) LaKind, J. S.; Naiman, J.; Verner, M. A.; Leveque, L.; Fenton, S. Per- and polyfluoroalkyl substances (PFAS) in breast milk and infant formula: A global issue. Environ Res 2023, 219, 115042. DOI: 10.1016/j.envres.2022.115042 From NLM Medline.

(13) Mogensen, U. B.; Grandjean, P.; Nielsen, F.; Weihe, P.; Budtz-Jorgensen, E. Breastfeeding as an Exposure Pathway for Perfluorinated Alkylates. Environ Sci Technol 2015, 49 (17), 10466–10473. DOI: 10.1021/acs.est.5b02237 From NLM Medline.

(14) Dietert, R. R. Developmental immunotoxicology: focus on health risks. Chem Res Toxicol 2009, 22 (1), 17–23. DOI: 10.1021/tx800198m.

(15) Hertz-Picciotto, I.; Park, H. Y.; Dostal, M.; Kocan, A.; Trnovec, T.; Sram, R. Prenatal exposures to persistent and non-persistent organic compounds and effects on immune system development. Basic Clin Pharmacol Toxicol 2008, 102 (2), 146–154. DOI: 10.1111/j.1742-7843.2007.00190.x From NLM Medline.

(16) Winans, B.; Humble, M. C.; Lawrence, B. P. Environmental toxicants and the developing immune system: a missing link in the global battle against infectious disease? Reprod Toxicol 2011, 31 (3), 327–336. DOI: 10.1016/j.reprotox.2010.09.004 From NLM Medline.

(17) Grandjean, P.; Andersen, E. W.; Budtz-Jorgensen, E.; Nielsen, F.; Molbak, K.; Weihe, P.; Heilmann, C. Serum vaccine antibody concentrations in children exposed to perfluorinated compounds. JAMA 2012, 307 (4), 391–397. DOI: 10.1001/jama.2011.2034 From NLM Medline.

(18) Granum, B.; Haug, L. S.; Namork, E.; Stolevik, S. B.; Thomsen, C.; Aaberge, I. S.; van Loveren, H.; Lovik, M.; Nygaard, U. C. Pre-natal exposure to perfluoroalkyl substances may be associated with altered vaccine antibody levels and immune-related health outcomes in early childhood. J Immunotoxicol 2013, 10 (4), 373–379. DOI: 10.3109/1547691X.2012.755580 From NLM Medline.

(19) Crawford, L.; Halperin, S. A.; Dzierlenga, M. W.; Skidmore, B.; Linakis, M. W.; Nakagawa, S.; Longnecker, M. P. Systematic review and meta-analysis of epidemiologic data on vaccine response in relation to exposure to five principal perfluoroalkyl substances. Environ Int 2023, 172, 107734. DOI: 10.1016/j.envint.2023.107734 From NLM Medline.

(20) Grandjean, P.; Heilmann, C.; Weihe, P.; Nielsen, F.; Mogensen, U. B.; Timmermann, A.; Budtz-Jorgensen, E. Estimated exposures to perfluorinated compounds in infancy predict attenuated vaccine antibody concentrations at age 5-years. J Immunotoxicol 2017, 14 (1), 188–195. DOI: 10.1080/1547691X.2017.1360968 From NLM Medline.

(21) Sigvaldsen, A.; Hojsager, F. D.; Paarup, H. M.; Beck, I. H.; Timmermann, C. A. G.; Boye, H.; Nielsen, F.; Halldorsson, T. I.; Nielsen, C.; Moller, S.;, et al. Early-life exposure to perfluoroalkyl substances and serum antibody concentrations towards common childhood vaccines in 18-month-old children in the Odense Child Cohort. Environ Res 2024, 242, 117814. DOI: 10.1016/j.envres.2023.117814 From NLM Medline.

(22) Hong, X.; Morgenlander, W. R.; Nadeau, K.; Wang, G.; Frischmeyer-Guerrerio, P. A.; Pearson, C.; Adams, W. G.; Ji, H.; Larman, H. B.; Wang, X. Maternal exposure to per- and polyfluoroalkyl substances and epitope level antibody response to vaccines against measles and rubella in children from the Boston birth cohort. Environ Int 2025, 198, 109433. DOI: 10.1016/j.envint.2025.109433 From NLM Medline.

(23) Lam, N.; Lee, Y.; Farber, D. L. A guide to adaptive immune memory. Nat Rev Immunol 2024, 24 (11), 810–829. DOI: 10.1038/s41577-024-01040-6 From NLM Medline.

(24) Van Loveren, H.; Van Amsterdam, J. G.; Vandebriel, R. J.; Kimman, T. G.; Rumke, H. C.; Steerenberg, P. S.; Vos, J. G. Vaccine-induced antibody responses as parameters of the influence of endogenous and environmental factors. Environ Health Perspect 2001, 109 (8), 757–764. DOI: 10.1289/ehp.01109757 From NLM Medline.

(25) Hikono, H.; Kohlmeier, J. E.; Ely, K. H.; Scott, I.; Roberts, A. D.; Blackman, M. A.; Woodland, D. L. T-cell memory and recall responses to respiratory virus infections. Immunol Rev 2006, 211, 119–132. DOI: 10.1111/j.0105-2896.2006.00385.x From NLM Medline.

(26) Ehrlich, V.; Bil, W.; Vandebriel, R.; Granum, B.; Luijten, M.; Lindeman, B.; Grandjean, P.; Kaiser, A. M.; Hauzenberger, I.; Hartmann, C.;, et al. Consideration of pathways for immunotoxicity of per- and polyfluoroalkyl substances (PFAS). Environ Health 2023, 22 (1), 19. DOI: 10.1186/s12940-022-00958-5 From NLM Medline.

(27) Melendez, D. C.; Laniewski, N.; Jusko, T. A.; Qiu, X.; Lawrence, B. P.; Rivera-Nunez, Z.; Brunner, J.; Best, M.; Macomber, A.; Leger, A.;, et al. In utero per - and polyfluoroalkyl substances (PFAS) exposure and changes in infant T helper cell development among UPSIDE-ECHO cohort participants. Environ Health Perspect 2025. DOI: 10.1289/EHP16726 From NLM Publisher.

(28) Antoniou, E.; Colnot, T.; Zeegers, M.; Dekant, W. Immunomodulation and exposure to per- and polyfluoroalkyl substances: an overview of the current evidence from animal and human studies. Arch Toxicol 2022, 96 (8), 2261–2285. DOI: 10.1007/s00204-022-03303-4 From NLM Medline.

(29) Dewitt, J.; Copeland, C.; Strynar, M.; Luebke, R. W. Perfluorooctanoic acid-induced immunomodulation in adult C57BL/6J or C57BL/6N female mice Environ Health Persp 2008, 116 (5), 644–650. DOI: doi: 10.1289/ehp.10896.

(30) Peden-Adams, M. M.; Keller, J. M.; Eudaly, J. G.; Berger, J.; Gilkeson, G. S.; Keil, D. E. Suppression of humoral immunity in mice following exposure to perfluorooctane sulfonate. Toxicol Sci 2008, 104, 144–154.

(31) Keil, D. E.; Mehlmann, T.; Butterworth, L.; Peden-Adams, M. M. Gestational Exposure to Perfluorooctane Sulfonate Suppresses Immune Function in B6C3F1 Mice. Toxicol Sci 2008, 103, 77–85.

(32) Torres, L.; Edko, A.; Limper, C.; Imbiakha, B.; Chang, S.; August, A. Effect of Perfluorooctanesulfonic acid (PFOS) on immune cell development and function in mice Immunol Lett 2021, 233, 31–41. DOI: doi: 10.1016/j.imlet.2021.03.006.

(33) Rockwell, C. E.; Turley, A. E.; Cheng, X.; Fields, P. E.; Klaassen, C. D. Persistent alterations in immune cell populations and function from a single dose of perfluorononanoic acid (PFNA) in C57Bl/6 mice. Food Chem Toxicol 2017, 100, 24–33. DOI: 10.1016/j.fct.2016.12.004 From NLM Medline.

(34) Ojo, A. F.; Peng, C.; Ng, J. C. Assessing the human health risks of per- and polyfluoroalkyl substances: A need for greater focus on their interactions as mixtures. J Hazard Mater 2021, 407, 124863. DOI: 10.1016/j.jhazmat.2020.124863 From NLM Medline.

(35) Wallis, D. J.; Barton, K. E.; Knappe, D. R. U.; Kotlarz, N.; McDonough, C. A.; Higgins, C. P.; Hoppin, J. A.; Adgate, J. L. Source apportionment of serum PFASs in two highly exposed communities. Sci Total Environ 2023, 855, 158842. DOI: 10.1016/j.scitotenv.2022.158842 From NLM Medline.

(36) Zhang, Y.; Mustieles, V.; Sun, Y.; Oulhote, Y.; Wang, Y. X.; Messerlian, C. Association between serum per- and polyfluoroalkyl substances concentrations and common cold among children and adolescents in the United States. Environ Int 2022, 164, 107239. DOI: 10.1016/j.envint.2022.107239 From NLM Medline.

(37) Padula, A. M.; Salihovic, S.; Zazara, D. E.; Diemert, A.; Arck, P. C. Prenatal per- and polyfluoroalkyl substances in relation to antibody titers and infections in childhood. Environ Res 2025, 270, 120976. DOI: 10.1016/j.envres.2025.120976 From NLM Medline.

(38) Landreth, K. S. Critical windows in development of the rodent immune system. Hum Exp Toxicol 2002, 21 (9-10), 493–498. DOI: 10.1191/0960327102ht287oa.

(39) Perez, F.; Nadal, M.; Navarro-Ortega, A.; Fabrega, F.; Domingo, J. L.; Barcelo, D.; Farre, M. Accumulation of perfluoroalkyl substances in human tissues. Environ Int 2013, 59, 354–362. DOI: 10.1016/j.envint.2013.06.004 From NLM Medline.

(40) Winans, B.; Nagari, A.; Chae, M.; Post, C. M.; Ko, C. I.; Puga, A.; Kraus, W. L.; Lawrence, B. P. Linking the aryl hydrocarbon receptor with altered DNA methylation patterns and developmentally induced aberrant antiviral CD8+ T cell responses. J Immunol 2015, 194 (9), 4446–4457. DOI: 10.4049/jimmunol.1402044.

(41) Warren, T. K.; Mitchell, K. A.; Lawrence, B. P. Exposure to 2,3,7,8-tetrachlorodibenzo-p-dioxin (TCDD) suppresses the humoral and cell-mediated immune responses to influenza A virus without affecting cytolytic activity in the lung. Toxicological sciences : an official journal of the Society of Toxicology 2000, 56 (1), 114–123.

(42) Boule, L. A.; Winans, B.; Lawrence, B. P. Effects of developmental activation of the AhR on CD4+ T-cell responses to influenza virus infection in adult mice. Environ Health Perspect 2014, 122 (11), 1201–1208. DOI: 10.1289/ehp.1408110.

(43) Barrett, T., Inglis, SC. Growth, purification, and titration of influenza viruses. In Virology: A Practical Approach, Mahy, B. Ed.; IRL Press, 1985; pp 119–150.

(44) Boule, L. A.; Burke, C. G.; Jin, G. B.; Lawrence, B. P. Aryl hydrocarbon receptor signaling modulates antiviral immune responses: ligand metabolism rather than chemical source is the stronger predictor of outcome. Sci Rep 2018, 8 (1), 1826. DOI: 10.1038/s41598-018-20197-4 From NLM Medline.

(45) Houser, C. L.; Lawrence, B. P. The Aryl Hydrocarbon Receptor Modulates T Follicular Helper Cell Responses to Influenza Virus Infection in Mice. J Immunol 2022, 208 (10), 2319–2330. DOI: 10.4049/jimmunol.2100936 From NLM Medline.

(46) Post, C. M.; McDonough, C.; Lawrence, B. P. Binary and quaternary mixtures of perfluoroalkyl substances (PFAS) differentially affect the immune response to influenza A virus infection. J Immunotoxicol 2024, 21 (1), 2340495. DOI: 10.1080/1547691X.2024.2340495 From NLM Medline.

(47) Dauner, J. G.; Pan, Y.; Hildesheim, A.; Kemp, T. J.; Porras, C.; Pinto, L. A. Development and application of a GuHCl-modified ELISA to measure the avidity of anti-HPV L1 VLP antibodies in vaccinated individuals. Mol Cell Probes 2012, 26 (2), 73–80. DOI: 10.1016/j.mcp.2012.01.002 From NLM Medline.

(48) Houser, C. L.; Fenner, K. N.; Lawrence, B. P. Timing influences the impact of aryl hydrocarbon receptor activation on the humoral immune response to respiratory viral infection. Toxicol Appl Pharmacol 2024, 489, 117010. DOI: 10.1016/j.taap.2024.117010 From NLM Medline.

(49) Livak, K. J.; Schmittgen, T. D. Analysis of relative gene expression data using real-time quantitative PCR and the 2(-Delta Delta C(T)) Method. Methods 2001, 25 (4), 402–408. DOI: 10.1006/meth.2001.1262 From NLM Medline.

(50) Rosen, M. B.; Lee, J. S.; Ren, H.; Vallanat, B.; Liu, J.; Waalkes, M. P.; Abbott, B. D.; Lau, C.; Corton, J. C. Toxicogenomic dissection of the perfluorooctanoic acid transcript profile in mouse liver: evidence for the involvement of nuclear receptors PPAR alpha and CAR. Toxicological sciences : an official journal of the Society of Toxicology 2008, 103 (1), 46–56. DOI: 10.1093/toxsci/kfn025 From NLM Medline.

(51) Guelfo, J. L.; Adamson, D. T. Evaluation of a national data set for insights into sources, composition, and concentrations of per- and polyfluoroalkyl substances (PFASs) in U.S. drinking water. Environ Pollut 2018, 236, 505–513. DOI: 10.1016/j.envpol.2018.01.066 From NLM Medline.

(52) USEPA, E. P. A. Per- and Polyfluoroalkyl Substances National Primary Drinking Water Regulation. In EPA-HQ-OW-2022-0114-0027, Agency, E. P., Ed.; 2023; pp 18638-18754.

(53) Dewitt, J. C.; Copeland, C. B.; Strynar, M. J.; Luebke, R. W. Perfluorooctanoic acid-induced immunomodulation in adult C57BL/6J or C57BL/6N female mice. Environ Health Perspect 2008, 116 (5), 644–650. DOI: 10.1289/ehp.10896 From NLM Medline.

(54) Evans, N.; Conley, J. M.; Cardon, M.; Hartig, P.; Medlock-Kakaley, E.; Gray, L. E., Jr. In vitro activity of a panel of per- and polyfluoroalkyl substances (PFAS), fatty acids, and pharmaceuticals in peroxisome proliferator-activated receptor (PPAR) alpha, PPAR gamma, and estrogen receptor assays. Toxicol Appl Pharmacol 2022, 449, 116136. DOI: 10.1016/j.taap.2022.116136 From NLM Medline.

(55) Rosen, M. B.; Das, K. P.; Rooney, J.; Abbott, B.; Lau, C.; Corton, J. C. PPARalpha-independent transcriptional targets of perfluoroalkyl acids revealed by transcript profiling. Toxicology 2017, 387, 95–107. DOI: 10.1016/j.tox.2017.05.013 From NLM Medline.

(56) Schlezinger, J. J.; Puckett, H.; Oliver, J.; Nielsen, G.; Heiger-Bernays, W.; Webster, T. F. Perfluorooctanoic acid activates multiple nuclear receptor pathways and skews expression of genes regulating cholesterol homeostasis in liver of humanized PPARalpha mice fed an American diet. Toxicol Appl Pharmacol 2020, 405, 115204. DOI: 10.1016/j.taap.2020.115204 From NLM Medline.

(57) Lawrence, B. P.; Roberts, A. D.; Neumiller, J. J.; Cundiff, J. A.; Woodland, D. L. Aryl hydrocarbon receptor activation impairs the priming but not the recall of influenza virus-specific CD8+ T cells in the lung. J Immunol 2006, 177 (9), 5819–5828. DOI: 10.4049/jimmunol.177.9.5819 From NLM Medline.

(58) Fink, A. L.; Engle, K.; Ursin, R. L.; Tang, W. Y.; Klein, S. L. Biological sex affects vaccine efficacy and protection against influenza in mice. Proc Natl Acad Sci U S A 2018, 115 (49), 12477–12482. DOI: 10.1073/pnas.1805268115 From NLM Medline.

(59) Boyden, A. W.; Legge, K. L.; Waldschmidt, T. J. Pulmonary infection with influenza A virus induces site-specific germinal center and T follicular helper cell responses. PLoS One 2012, 7 (7), e40733. DOI: 10.1371/journal.pone.0040733 From NLM Medline.

(60) Chapman, T. J.; Castrucci, M. R.; Padrick, R. C.; Bradley, L. M.; Topham, D. J. Antigen-specific and non-specific CD4+ T cell recruitment and proliferation during influenza infection. Virology 2005, 340 (2), 296–306. DOI: 10.1016/j.virol.2005.06.023 From NLM Medline.

(61) Swain, S. L.; McKinstry, K. K.; Strutt, T. M. Expanding roles for CD4(+) T cells in immunity to viruses. Nat Rev Immunol 2012, 12 (2), 136–148. DOI: 10.1038/nri3152 From NLM Medline.

(62) Crotty, S. Follicular helper CD4 T cells (TFH). Annu Rev Immunol 2011, 29, 621–663. DOI: 10.1146/annurev-immunol-031210-101400 From NLM Medline.

(63) Belanger, S.; Crotty, S. Dances with cytokines, featuring TFH cells, IL-21, IL-4 and B cells. Nat Immunol 2016, 17 (10), 1135-1136. DOI: 10.1038/ni.3561 From NLM Medline.

(64) Krautler, N. J.; Suan, D.; Butt, D.; Bourne, K.; Hermes, J. R.; Chan, T. D.; Sundling, C.; Kaplan, W.; Schofield, P.; Jackson, J.;, et al. Differentiation of germinal center B cells into plasma cells is initiated by high-affinity antigen and completed by Tfh cells. J Exp Med 2017, 214 (5), 1259–1267. DOI: 10.1084/jem.20161533 From NLM Medline.

(65) Barton, K. E.; Zell-Baran, L. M.; DeWitt, J. C.; Brindley, S.; McDonough, C. A.; Higgins, C. P.; Adgate, J. L.; Starling, A. P. Cross-sectional associations between serum PFASs and inflammatory biomarkers in a population exposed to AFFF-contaminated drinking water. Int J Hyg Environ Health 2022, 240, 113905. DOI: 10.1016/j.ijheh.2021.113905 From NLM Medline.

(66) Bline, A. P.; DeWitt, J. C.; Kwiatkowski, C. F.; Pelch, K. E.; Reade, A.; Varshavsky, J. R. Public Health Risks of PFAS-Related Immunotoxicity Are Real. Curr Environ Health Rep 2024, 11 (2), 118–127. DOI: 10.1007/s40572-024-00441-y From NLM Medline.

(67) Bulka, C. M.; Avula, V.; Fry, R. C. Associations of exposure to perfluoroalkyl substances individually and in mixtures with persistent infections: Recent findings from NHANES 1999-2016. Environ Pollut 2021, 275, 116619. DOI: 10.1016/j.envpol.2021.116619 From NLM Medline.

(68) Lyrou, M.; Karampas, G.; Metallinou, D.; Athanasiadou, C. R.; Gourounti, K.; Jotautis, V.; Georgakopoulou, V. E.; Spandidos, D. A.; Sarantaki, A. Association between prenatal exposure to perfluoroalkyl and polyfluoroalkyl substances and the incidence of infant and childhood respiratory infections: A systematic review. Med Int (Lond) 2025, 5 (5), 51. DOI: 10.3892/mi.2025.250 From NLM PubMed-not-MEDLINE.

(69) Nutt, S. L.; Hodgkin, P. D.; Tarlinton, D. M.; Corcoran, L. M. The generation of antibody-secreting plasma cells. Nat Rev Immunol 2015, 15 (3), 160–171. DOI: 10.1038/nri3795 From NLM Medline.

(70) Rudzanova, B.; Thon, V.; Vespalcova, H.; Martyniuk, C. J.; Piler, P.; Zvonar, M.; Klanova, J.; Blaha, L.; Adamovsky, O. Altered Transcriptome Response in PBMCs of Czech Adults Linked to Multiple PFAS Exposure: B Cell Development as a Target of PFAS Immunotoxicity. Environ Sci Technol 2024, 58 (1), 90–98. DOI: 10.1021/acs.est.3c05109 From NLM Medline.

(71) Zhang, J.; Chen, J.; Kang, R.; Dai, C.; Xie, B.; Li, K.; Sun, R.; Zheng, X.; Tan, L.; Xu, X.;, et al. Tfh cell-mediated impairment of rotavirus-specific antibodies by PAH and PFAS co-exposure in seropositive children. J Hazard Mater 2025, 497, 139630. DOI: 10.1016/j.jhazmat.2025.139630 From NLM Medline.

(72) Taylor, K. D.; Woodlief, T. L.; Ahmed, A.; Hu, Q.; Duncker, P. C.; DeWitt, J. C. Quantifying the impact of PFOA exposure on B-cell development and antibody production. Toxicological sciences : an official journal of the Society of Toxicology 2023, 194 (1), 101–108. DOI: 10.1093/toxsci/kfad043 From NLM Medline.

(73) DeWitt, J. C.; Williams, W. C.; Creech, N. J.; Luebke, R. W. Suppression of antigen-specific antibody responses in mice exposed to perfluorooctanoic acid: Role of PPARalpha and T- and B-cell targeting. J Immunotoxicol 2016, 13 (1), 38–45. DOI: 10.3109/1547691X.2014.996682 From NLM Medline.

(74) Peden-Adams, M. M.; Keller, J. M.; Eudaly, J. G.; Berger, J.; Gilkeson, G. S.; Keil, D. E. Suppression of humoral immunity in mice following exposure to perfluorooctane sulfonate. Toxicological sciences : an official journal of the Society of Toxicology 2008, 104 (1), 144–154. DOI: 10.1093/toxsci/kfn059 From NLM Medline.

(75) Keil, D. E.; Mehlmann, T.; Butterworth, L.; Peden-Adams, M. M. Gestational exposure to perfluorooctane sulfonate suppresses immune function in B6C3F1 mice. Toxicological sciences : an official journal of the Society of Toxicology 2008, 103 (1), 77–85. DOI: 10.1093/toxsci/kfn015 From NLM Medline.

(76) Rockwell, C. E.; Turley, A. E.; Cheng, X.; Fields, P. E.; Klaassen, C. D. Acute Immunotoxic Effects of Perfluorononanoic Acid (PFNA) in C57BL/6 Mice. Clin Exp Pharmacol 2013, *Suppl* 4. DOI: 10.4172/2161-1459.S4-002 From NLM PubMed-not-MEDLINE.

(77) Lorenzo, M. E.; Hodgson, A.; Robinson, D. P.; Kaplan, J. B.; Pekosz, A.; Klein, S. L. Antibody responses and cross protection against lethal influenza A viruses differ between the sexes in C57BL/6 mice. Vaccine 2011, 29 (49), 9246–9255. DOI: 10.1016/j.vaccine.2011.09.110 From NLM Medline.

(78) Zazara, D. E.; Arck, P. C. Developmental origin and sex-specific risk for infections and immune diseases later in life. Semin Immunopathol 2019, 41 (2), 137–151. DOI: 10.1007/s00281-018-0713-x From NLM Medline.

(79) Corsini, E.; Luebke, R. W.; Germolec, D. R.; DeWitt, J. C. Perfluorinated compounds: emerging POPs with potential immunotoxicity. Toxicol Lett 2014, 230 (2), 263–270. DOI: 10.1016/j.toxlet.2014.01.038 From NLM Medline.

(80) Abbott, B. D. Review of the expression of peroxisome proliferator-activated receptors alpha (PPAR alpha), beta (PPAR beta), and gamma (PPAR gamma) in rodent and human development. Reprod Toxicol 2009, 27 (3-4), 246–257. DOI: 10.1016/j.reprotox.2008.10.001 From NLM Medline.

(81) Christofides, A.; Konstantinidou, E.; Jani, C.; Boussiotis, V. A. The role of peroxisome proliferator-activated receptors (PPAR) in immune responses. Metabolism 2021, 114, 154338. DOI: 10.1016/j.metabol.2020.154338 From NLM Medline.

(82) Szilagyi, J. T.; Avula, V.; Fry, R. C. Perfluoroalkyl Substances (PFAS) and Their Effects on the Placenta, Pregnancy, and Child Development: a Potential Mechanistic Role for Placental Peroxisome Proliferator-Activated Receptors (PPARs). Curr Environ Health Rep 2020, 7 (3), 222–230. DOI: 10.1007/s40572-020-00279-0 From NLM Medline.

(83) Park, H. J.; Choi, J. M. Sex-specific regulation of immune responses by PPARs. Exp Mol Med 2017, 49 (8), e364. DOI: 10.1038/emm.2017.102 From NLM Medline.

(84) Park, H. J.; Park, H. S.; Lee, J. U.; Bothwell, A. L.; Choi, J. M. Gender-specific differences in PPARgamma regulation of follicular helper T cell responses with estrogen. Sci Rep 2016, 6, 28495. DOI: 10.1038/srep28495 From NLM Medline.

(85) Christodoulou, M.; Moysidou, E.; Lioulios, G.; Stai, S.; Lazarou, C.; Xochelli, A.; Fylaktou, A.; Stangou, M. T-Follicular Helper Cells and Their Role in Autoimmune Diseases. Life (Basel) 2025, 15 (4). DOI: 10.3390/life15040666 From NLM PubMed-not-MEDLINE.

