## Supplemental information for "Developmental exposure to a PFAS mixture impairs the anamnestic response to influenza A virus infection in mice"

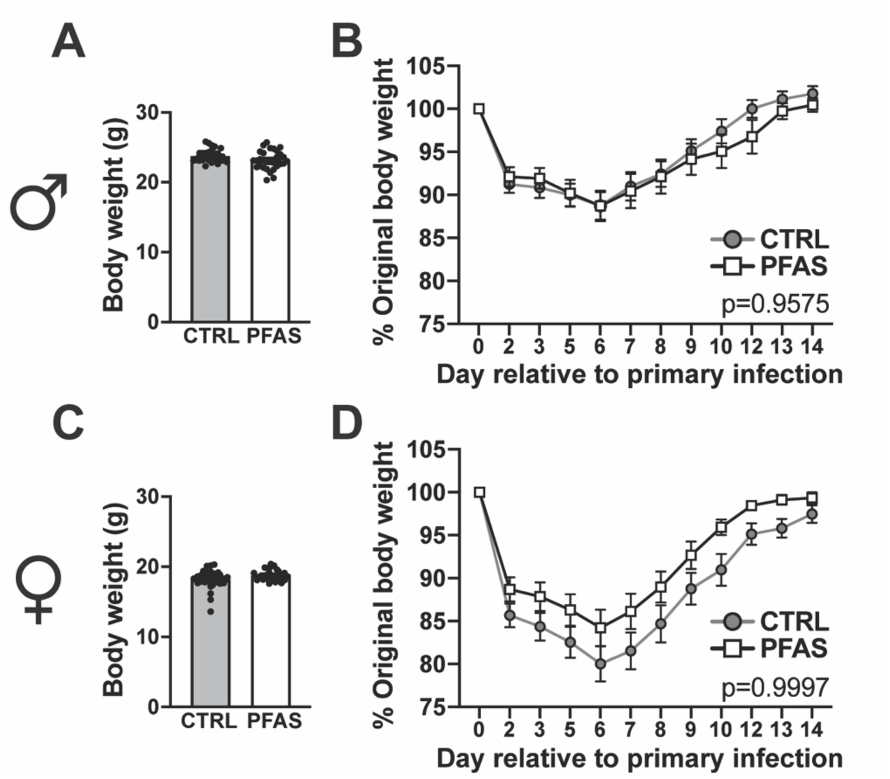

**Figure S1. Developmental exposure to the PFAS mixture did not affect body weight change during primary IAV infection. (A)** Mean body weight of male offspring on the day of primary IAV infection (day 0); n=33 CTRL, 30 PFAS. **(B)** Mean body weight (expressed as a percentage of day 0 body weight) of male offspring after primary IAV infection. **(C)** Mean body weight of female offspring on day 0; n=30 CTRL, 28 PFAS. **(D)** Mean body weight of female offspring during primary IAV infection (expressed as a percentage of day 0 body weight). Data are expressed as mean ± SEM. **(B, D)** Body weight data over time were analyzed by two-way ANOVA followed by Tukey's HSD post hoc test. The p-value for the interaction between treatment and day relative to infection is indicated on the graph.

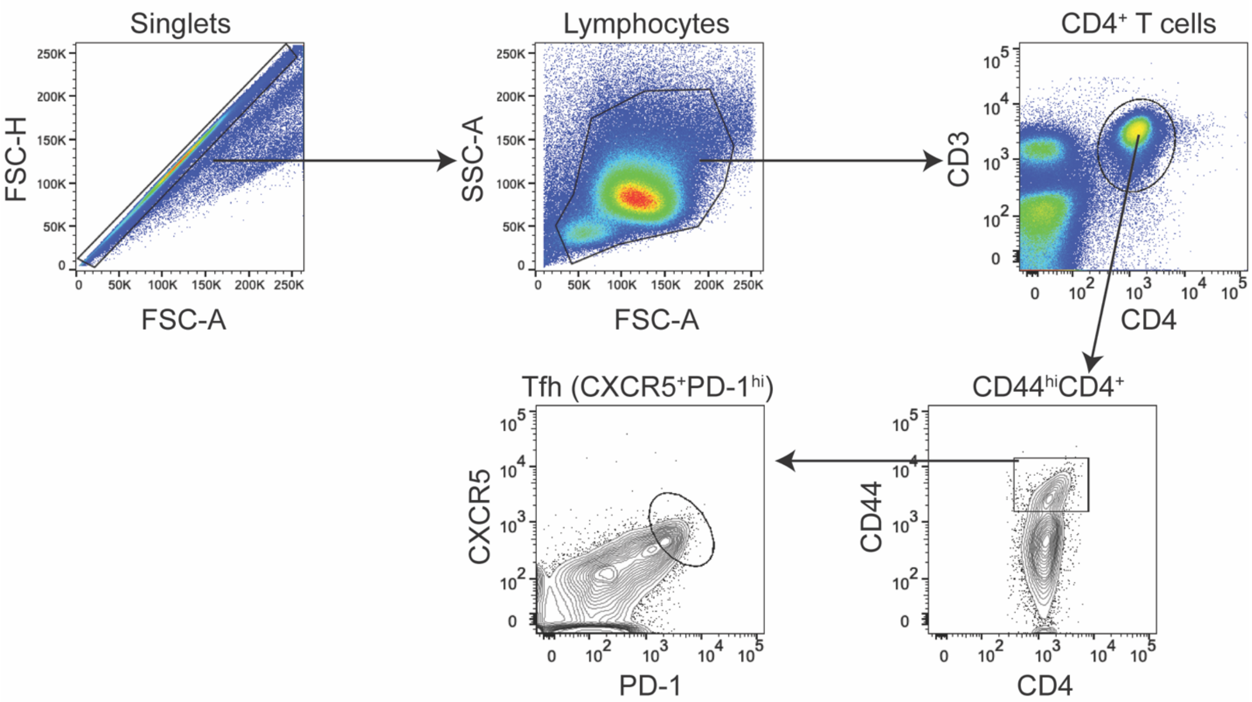

**Figure S2. Tfh cell gating strategy.** Single cell suspensions of MLN cells were subjected to analysis by flow cytometry. Doublets, debris and nonviable cells were excluded by gating on singlets. Side scatter area (SSC-A) and forward scatter area (FSC-A) were used to gate broadly on the total lymphocyte population. Co-expression of CD3 and CD4 was used to identify CD4+ T cells. After gating on CD44hiCD4+ T cells, Tfh cells ($CD4^{+}CD44^{hi}CXCR5^{+}PD\text{-}1^{hi}$) were identified based on expression of CXCR5 and PD-1.

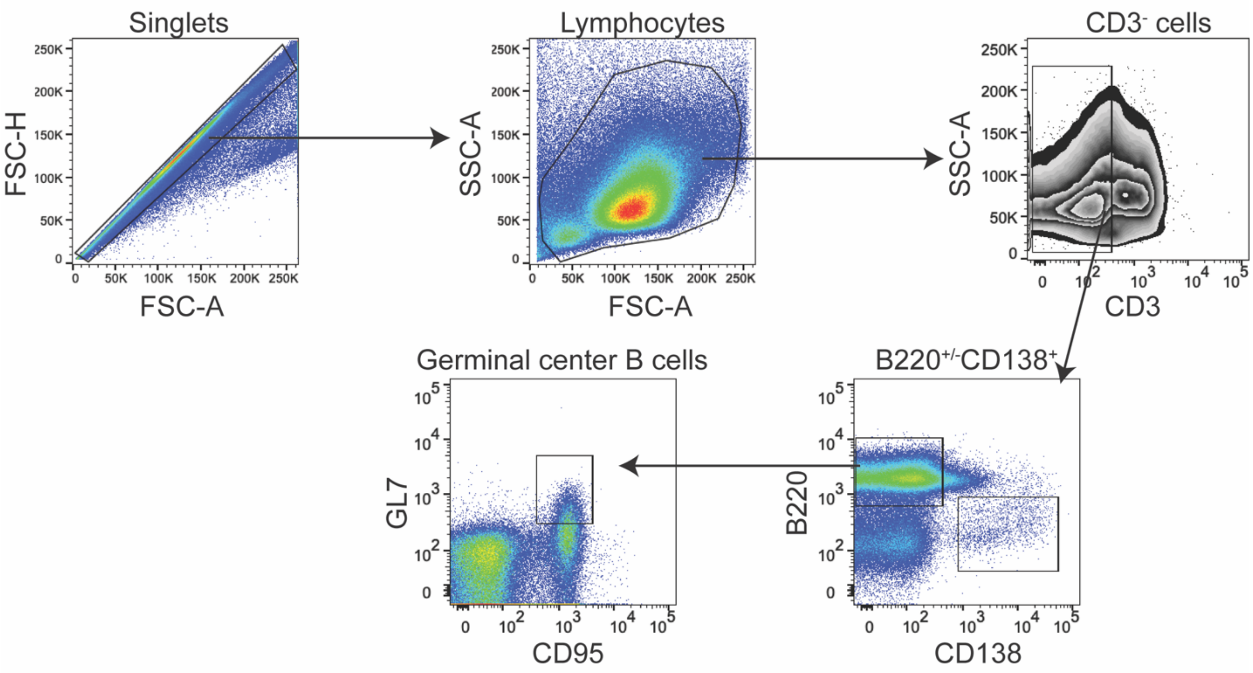

**Figure S3. Gating strategy to identify germinal center B cells.** Single cell suspensions of MLN cells were subjected to analysis by flow cytometry. Doublets and nonviable cells were excluded by gating on singlets. Side scatter area (SSC-A) and forward scatter area (FSC-A) were used to identify the total lymphocyte population. T cells were excluded by gating on the CD3 negative population. After gating on B220+CD138- cells, germinal center B cells ($CD3^{-}B220^{+}CD95^{+}GL7^{+}$) were identified based on coexpression of GL7 and CD95.

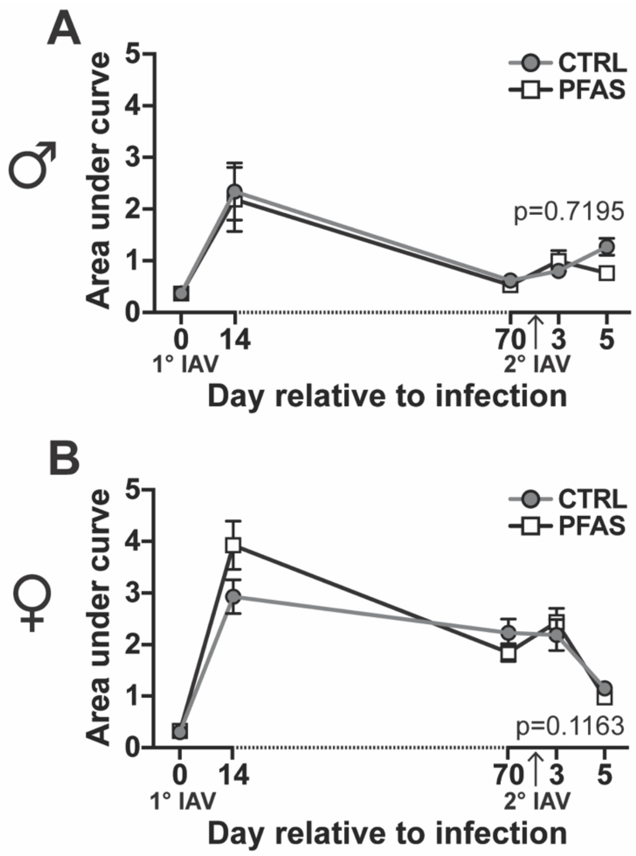

**Figure S4. Developmental exposure to the PFAS mixture does not affect IAV-specific IgM levels.** Male and female offspring of control and PFAS-treated dams were infected with IAV as depicted in **Figure 3A**. Serum was collected at the indicated time points before and after first and second infection. The first infection was on day 0, and the second infection was on day 72 (upward arrow on x-axes). The relative levels of circulating IAV-specific IgM were measured by ELISA, using serially diluted serum. Area under curve (AUC) analysis of each dilution series was performed. Graphs depict mean AUC values of IAV-specific IgM levels throughout primary and secondary IAV infection in **(A)** male offspring (n=3-6 mice per time point) and **(B)** female offspring (n=4-6 mice per time point). Data are presented as mean ± SEM and were compared by two-way ANOVA followed by Tukey's HSD post hoc test. The p-value for the interaction between treatment and day relative to infection is listed on each graph. A full list of AUC values and p-values is provided in **Table S9**.

**Table S1.** Flow cytometry antibodies and reagents.

| Antibody/Reagent[a] | Clone | Company | Catalog No. | Amount /2x$10^6$ cells |
|---|---|---|---|---|
| CD3ε PE | 145-2C11 | BD Biosciences | 553064 | 0.06 µg |
| CD3ε PE-CF594 | 145-2C11 | BD Biosciences | 562286 | 0.25 µg |
| CD4 BV650 | RM4-5 | BD Biosciences | 563747 | 0.06 µg |
| CD8α APC-Cy7 | 53-6.7 | BD Biosciences | 557654 | 0.25 µg |
| CD19 BV421 | 1D3 | BD Biosciences | 562701 | 0.06 µg |
| CD19 BV605 | 1D3 | BD Biosciences | 563148 | 0.06 µg |
| CD19 BV711 | 1D3 | BD Biosciences | 563157 | 0.06 µg |
| CD19 PE | 1D3 | BD Biosciences | 553786 | 0.06 µg |
| CD25 APC | PC61 | BD Biosciences | 557192 | 0.50 µg |
| CD44 BV605 | IM7 | BD Biosciences | 563058 | 0.50 µg |
| CD45R/B220 AF488 | RA3-6B2 | BD Biosciences | 557669 | 0.06 µg |
| CD45R/B220 AF700 | RA3-6B2 | BD Biosciences | 557957 | 0.125 µg |
| CD45R/B220 APC | RA3-6B2 | BD Biosciences | 553092 | 0.06 µg |
| CD45R/B220 BV605 | RA3-6B2 | BD Biosciences | 563708 | 0.06 µg |
| CD45R/B220 eF450 | RA3-6B2 | Invitrogen | 48-0452-82 | 0.125 µg |
| CD45R/B220 FITC | RA3-6B2 | Invitrogen | 11-0452-82 | 0.16 µg |
| CD45R/B220 PE-Cy7 | RA3-6B2 | BD Biosciences | 552772 | 0.125 µg |
| CD95 PE-Cy7 | Jo2 | BD Biosciences | 557653 | 0.50 µg |
| CD138 BV711 | 281-2 | BD Biosciences | 563193 | 0.06 µg |
| CXCR5 (CD185) Biotin | 2G8 | BD Biosciences | 551960 | 1.25 µg |
| PD-1 (CD279) BV421 | J43 | BD Biosciences | 562584 | 0.25 µg |
| FOXP3 AF700 | FJK-16s | Invitrogen | 56-5773-82 | 0.25 µg |
| GATA3 AF488 | L50-823 | BD Biosciences | 560163 | 10.0 µl |
| GL7 eF450 | GL7 | Invitrogen | 48-5902-82 | 0.25 µg |
| RORγt PE | Q31-378 | BD Biosciences | 562607 | 0.20 µg |
| Streptavidin PE | NA | BD Biosciences | 554061 | 0.16 µg |
| Tbet PE-Cy7 | 4B10 | Invitrogen | 25-5825-82 | 0.25 µg |
| CD16/32 unconjugated | 93 | Invitrogen | 14-0161-86 | NA |
| BD Horizon Brilliant Stain Buffer | NA | BD Biosciences | 566349 | 50.0 µl |
| FoxP3 Transcription Factor Staining Buffer Set | NA | Invitrogen | 00-5523-00 | NA |

Abbreviations used: AF: Alexa Fluor, APC: allophycocyanin, BV: Brilliant Violet, CF: propriety annotation, Cy: cyanine, eF: eFluor, FITC: fluorescein isothiocyanate, PE: phycoerythrin, NA: not applicable
[a]Antibody information includes antigen and fluorochrome.

**Table S2**. Pregnancy outcomes in CTRL and PFAS-treated dams.

| | CTRL water | 1x PFAS water | 0.1x PFAS water | 0.01x PFAS water | 0.0001x PFAS water | p-value[a] |
|---|---|---|---|---|---|---|
| **Dam n** | 30[b] | 3 | 30[b] | 3 | 3 | NA |
| **% Not pregnant** | 6.7 | 33.3 | 23.3 | 0 | 33.3 | NA |
| **Time to parturition (d)** | 18.75 ± 0.45 | 19.0 ± 0 | 18.78 ± 0.74 | 18.5 ± 0.71 | 19.0 ± 1.41 | 0.9393 |
| **Litter size** [c] | 5.68 ± 1.86 | 3.0 ± 1.41 | 5.81 ± 1.57 | 5.67 ± 0.58 | 5.5 ± 3.54 | 0.3291 |
| **Offspring sex ratio (M: F)**[d] | 1.66 ± 1.48 | 0.5 ± 0.71 | 1.5 ± 1.14 | 0.33 ± 0.15 | 1.75 ± 1.77 | 0.4170 |

NA: Not applicable
[a]p-value calculated by one-way ANOVA.
[b]Total number of dams from three independent exposure groups.
[c]Litter size: number of live offspring at weaning (PND21).
[d]Sex ratio of live offspring at weaning.

**Table S3.** Offspring body weight, relative spleen and liver weight, and relative *Acox1* expression at weaning.

| Male offspring | | | | | | | | | | |
|---|---|---|---|---|---|---|---|---|---|---|
| | **Control water** | **1x PFAS water[a]** | | **0.1x PFAS water[a]** | | **0.01x PFAS water[a]** | | **0.0001x PFAS water[a]** | | **One-way ANOVA p-value** |
| | **Mean ± SEM** | **Mean ± SEM** | **p-value[b]** | **Mean ± SEM** | **p-value[b]** | **Mean ± SEM** | **p-value[b]** | **Mean ± SEM** | **p-value[b]** | |
| **Body weight (g)** | 8.58 ± 0.18 | 3.50 ± 0.5 | <0.0001 | 8.75 ± 0.23 | 0.9742 | 8.00 ± 0.41 | 0.9034 | 7.86 ± 0.26 | 0.6227 | <0.0001 |
| **Spleen weight[c]** | 0.70 ± 0.05 | ND | NA | 0.52 ± 0.04 | 0.0850 | 0.65[d] | 0.9865 | 0.53 ± 0.06 | 0.1227 | 0.0701 |
| **Liver weight[c]** | 4.11 ± 0.30 | 4.48 ± 1.76 | 0.9902 | 5.78 ± 0.36 | 0.0086 | 4.03 ± 0.40 | 0.9999 | 4.28 ± 0.16 | 0.9968 | 0.0127 |
| ***Acox1* fold-change** | 1.00 ± 0 | 6.49[d] | <0.0001 | 2.17 ± 0.25 | 0.0059 | 1.83 ± 0.27 | 0.0992 | 1.06 ± 0.13 | 0.9997 | <0.0001 |
| **Female offspring** | | | | | | | | | | |
| | **Control water** | **1x PFAS water[a]** | | **0.1x PFAS water[a]** | | **0.01x PFAS water[a]** | | **0.0001x PFAS water[a]** | | **One-way ANOVA p-value** |
| | **Mean ± SEM** | **Mean ± SEM** | **p-value[b]** | **Mean ± SEM** | **p-value[b]** | **Mean ± SEM** | **p-value[b]** | **Mean ± SEM** | **p-value[b]** | |
| **Body weight (g)** | 7.56 ± 0.21 | 3.25 ± 0.25 | <0.0001 | 8.29 ± 0.17 | 0.0623 | 8.31 ± 0.17 | 0.2875 | 7.25 ± 0.48 | 0.9878 | <0.0001 |
| **Spleen weight[c]** | 0.70 ± 0.02 | ND | NA | 0.56 ± 0.11 | 0.7085 | 0.68 ± 0.10 | 0.9986 | 0.57 ± 0.15 | 0.7510 | 0.6138 |
| **Liver weight[c]** | 3.99 ± 0.27 | 7.15 ± 0.27 | <0.0001 | 5.45 ± 0.42 | 0.0238 | 4.68 ± 0.12 | 0.1758 | 4.34 ± 0.15 | 0.9409 | <0.0001 |
| ***Acox1* fold-change** | 1.00 ± 0 | 7.33 ± 0.54 | <0.0001 | 3.92 ± 0.43 | <0.0001 | 2.45 ± 0.19 | 0.0059 | 2.06 ± 0.37 | 0.2617 | <0.0001 |

ND: not determined; NA: not applicable

[a]Denotes PFAS mixture concentration of water consumed by the dam.

[b]p-value of PFAS vs. control calculated by Tukey's HSD post hoc test.

[c]Organ weight is expressed as a percentage of body weight.

[d]n=1

**Table S4.** MLN cellularity and number of CD4+ T cells, CD8+ T cells, and CD19+ B cells in the MLN.

| Cell type | Day relative to primary infection | Male CTRL[a] | p-value[b] | Male PFAS[a] | p-value[b] | p-value[c] | Female CTRL[a] | p-value[b] | Female PFAS[a] | p-value[b] | p-value[c] | p-value CTRL male vs. CTRL female[d] | p-value PFAS male vs. PFAS female[d] |
|---|---|---|---|---|---|---|---|---|---|---|---|---|---|
| **Total MLN cells** | **0** | 1867500 ± 710624 | 1.0000 | 1112350 ± 321578 | 1.0000 | 0.9999 | 1476800 ± 318922 | 1.0000 | 2197500 ± 642379 | 1.0000 | 1.0000 | 1.0000 | 0.9941 |
| | **9** | 4772500 ± 1004311 | 0.1529 | 3311667 ± 323166 | 0.0232 | 0.9629 | 4050667 ± 939664 | 0.2629 | 3893333 ± 445643 | 0.3739 | 1.0000 | 1.0000 | 0.9999 |
| | **14** | 3945000 ± 1128615 | 0.3323 | 3608500 ± 735394 | 0.0090 | 1.0000 | 4991000 ± 1073385 | 0.0743 | 3656333 ± 739908 | 0.5084 | 0.9918 | 0.9994 | 1.0000 |
| | **70** | 1789500 ± 406461 | 1.0000 | 1644750 ± 208581 | 0.9316 | 1.0000 | 2134750 ± 496605 | 0.9910 | 3294333 ± 1087533 | 0.8381 | 0.9998 | 1.0000 | 0.9260 |
| | **75 (3)[e]** | 3886500 ± 903355 | 0.3121 | 2948600 ± 623925 | 0.0834 | 0.9970 | 6231667 ± 1452335 | 0.0100 | 5618000 ± 938741 | 0.0207 | 1.0000 | 0.6848 | 0.1455 |
| | **77 (5)[e]** | 2984000 ± 393228 | 0.8019 | 2343333 ± 325911 | 0.3158 | 0.9999 | 5561667 ± 784875 | 0.0307 | 4222167 ± 684682 | 0.2287 | 0.9916 | 0.5943 | 0.4912 |
| **CD8+** | **0** | 360722 ± 144926 | 1.0000 | 143423 ± 51025 | 1.0000 | 0.8504 | 285825 ± 65686 | 1.0000 | 430858 ± 132368 | 1.0000 | 0.9991 | 1.0000 | 0.7158 |
| | **9** | 661975 ± 137182 | 0.1910 | 489977 ± 57592 | 0.0070 | 0.9299 | 609079 ± 136256 | 0.2726 | 678653 ± 80218 | 0.4560 | 1.0000 | 1.0000 | 0.8966 |
| | **70** | 244003 ± 53220 | 0.8105 | 226088 ± 21478 | 0.8409 | 1.0000 | 357614 ± 60406 | 0.9898 | 497349 ± 164897 | 0.9926 | 0.9998 | 0.9996 | 0.8426 |
| | **75 (3)[e]** | 434092 ± 102362 | 0.9439 | 363995 ± 83623 | 0.1066 | 0.9923 | 688129 ± 152840 | 0.1298 | 698805 ± 175392 | 0.4229 | 1.0000 | 0.7614 | 0.2312 |
| | **77 (5)[e]** | 381423 ± 40008 | 0.9995 | 300906 ± 69507 | 0.3223 | 0.9990 | 700377 ± 154769 | 0.1146 | 566070 ± 100793 | 0.8564 | 0.9984 | 0.5151 | 0.5631 |
| **CD4+** | **0** | 4561167 ± 174123 | 1.0000 | 198243 ± 65924 | 1.0000 | 0.6464 | 402676 ± 101237 | 1.0000 | 592979 ± 200123 | 1.0000 | 0.9969 | 1.0000 | 0.5389 |
| | **9** | 697490 ± 130447 | 0.2823 | 579858 ± 76609 | 0.0059 | 0.9927 | 669996 ± 141210 | 0.4589 | 718358 ± 79793 | 0.9374 | 1.0000 | 1.0000 | 0.9955 |
| | **70** | 260483 ± 61136 | 0.3757 | 250794 ± 21458 | 0.9697 | 1.0000 | 389264 ± 72974 | 1.0000 | 546402 ± 179299 | 0.9991 | 0.9998 | 0.9990 | 0.9010 |
| | **75 (3)[e]** | 386293 ± 68680 | 0.9359 | 370794 ± 78648 | 0.3094 | 1.0000 | 670531 ± 141486 | 0.4572 | 701768 ± 212590 | 0.9657 | 1.0000 | 0.6227 | 0.5194 |
| | **77 (5)[e]** | 400782 ± 41027 | 0.9728 | 329530 ± 70961 | 0.5322 | 0.9995 | 784409 ± 163086 | 0.1814 | 630107 ± 121264 | 0.9993 | 0.9980 | 0.2552 | 0.6317 |
| **CD19+** | **0** | 586397 ± 246145 | 1.0000 | 664613 ± 141992 | 1.0000 | 1.0000 | 517298 ± 98559 | 1.0000 | 616425 ± 185709 | 1.0000 | 1.0000 | 1.0000 | 1.0000 |
| | **14** | 1821620 ± 683395 | 0.2539 | 1739888 ± 442303 | 0.0845 | 1.0000 | 2422152 ± 613993 | 0.0519 | 1434250 ± 384354 | 0.3718 | 0.8458 | 0.9961 | 0.9996 |
| | **70** | 723146 ± 204418 | 0.9987 | 750840 ± 102122 | 0.9992 | 1.0000 | 833108 ± 169345 | 0.9766 | 1340643 ± 414023 | 0.6285 | 0.9996 | 1.0000 | 0.9923 |
| | **75 (3)[e]** | 2142920 ± 542536 | 0.1008 | 1593219 ± 382384 | 0.1775 | 0.9919 | 3301858 ± 803235 | 0.0033 | 2842366 ± 402802 | 0.0024 | 0.9993 | 0.7415 | 0.3064 |
| | **77 (5)[e]** | 1625850 ± 259687 | 0.3634 | 1258852 ± 157825 | 0.4892 | 0.9994 | 2955668 ± 360366 | 0.0102 | 2170793 ± 445275 | 0.0279 | 0.9566 | 0.5716 | 0.6118 |

[a]Mean number of cells ± SEM.
[b]p-value calculated by Dunnett's post-hoc test (day relative to primary infection compared to day 0 within sex/developmental exposure group).
[c]p-value calculated by Tukey HSD post-hoc test (PFAS vs. CTRL within sex on each day relative to primary infection).
[d]p-value calculated by Tukey HSD post-hoc test (male vs. female within developmental exposure group on each day relative to primary infection).
[e]Number in parenthesis () indicates day relative to secondary infection.

**Table S5.** Percentage of CD4+ T cells, CD8+ T cells, and CD19+ B cells in the MLN.

| Cell type | Day relative to primary infection | Male CTRL[a] | p-value[b] | Male PFAS[a] | p-value[b] | p-value[c] | Female CTRL[a] | p-value[b] | Female PFAS[a] | p-value[b] | p-value[c] | p-value CTRL male vs. CTRL female[d] | p-value PFAS male vs. PFAS female[d] |
|---|---|---|---|---|---|---|---|---|---|---|---|---|---|
| CD8+ | **0** | 18.38 ± 1.34 | 1.0000 | 10.90 ± 2.98 | 1.0000 | 0.0141 | 19.10 ± 1.05 | 1.0000 | 18.80 ± 1.47 | 1.0000 | 1.0000 | 1.0000 | 0.0254 |
| | **9** | 13.95 ± 0.44 | 0.0460 | 14.68 ± 0.53 | 0.2011 | 1.0000 | 15.60 ± 1.16 | 0.1443 | 17.42 ± 0.73 | 0.8727 | 0.9823 | 0.9913 | 0.8696 |
| | **70** | 13.68 ± 0.97 | 0.0181 | 14.00 ± 0.94 | 0.4226 | 1.0000 | 17.35 ± 1.14 | 0.7460 | 15.07 ± 0.09 | 0.3117 | 0.9875 | 0.4681 | 1.0000 |
| | **75 (3)[e]** | 11.40 ± 1.03 | 0.0003 | 10.50 ± 1.05 | 0.9984 | 1.0000 | 11.39 ± 1.24 | 0.0005 | 12.30 ± 2.15 | 0.0142 | 0.9999 | 1.0000 | 0.9975 |
| | **77 (5)[e]** | 13.11 ± 0.78 | 0.0060 | 12.05 ± 1.21 | 0.9323 | 0.9995 | 12.23 ± 1.21 | 0.0017 | 13.17 ± 1.07 | 0.0283 | 0.9999 | 0.9998 | 0.9997 |
| CD4+ | **0** | 23.78 ± 1.64 | 1.0000 | 15.13 ± 3.81 | 1.0000 | 0.0161 | 26.50 ± 1.36 | 1.0000 | 25.28 ± 2.56 | 1.0000 | 1.0000 | 0.9545 | 0.0231 |
| | **9** | 15.28 ± 1.71 | 0.0007 | 17.25 ± 0.82 | 0.7700 | 0.9940 | 17.75 ± 1.69 | 0.0022 | 18.63 ± 0.87 | 0.0638 | 1.0000 | 0.9683 | 0.9998 |
| | **70** | 14.32 ± 0.89 | <.0001 | 15.53 ± 0.78 | 0.9995 | 0.9999 | 18.75 ± 1.16 | 0.0143 | 16.53 ± 0.41 | 0.0358 | 0.9988 | 0.5013 | 1.0000 |
| | **75 (3)[e]** | 11.24 ± 1.08 | <.0001 | 11.17 ± 1.33 | 0.2991 | 1.0000 | 11.53 ± 1.91 | <.0001 | 12.27 ± 2.96 | 0.0005 | 1.0000 | 1.0000 | 1.0000 |
| | **77 (5)[e]** | 13.74 ± 0.76 | <.0001 | 13.31 ± 1.11 | 0.8449 | 1.0000 | 13.73 ± 1.15 | <.0001 | 14.37 ± 1.13 | 0.0019 | 1.0000 | 1.0000 | 1.0000 |
| CD19+ | **0** | 29.85 ± 1.61 | 1.0000 | 36.15 ± 1.68 | 1.0000 | 0.9817 | 35.74 ± 1.33 | 1.0000 | 31.76 ± 2.26 | 1.0000 | 0.9977 | 0.9803 | 0.9974 |
| | **14** | 41.20 ± 6.83 | 0.2039 | 45.40 ± 3.57 | 0.0638 | 0.9956 | 46.87 ± 2.81 | 0.0103 | 39.35 ± 4.68 | 0.5007 | 0.7809 | 0.9595 | 0.9374 |
| | **70** | 37.47 ± 4.26 | 0.5118 | 45.55 ± 2.15 | 0.0912 | 0.8629 | 38.18 ± 2.29 | 0.8701 | 41.70 ± 2.34 | 0.4305 | 0.9996 | 1.0000 | 0.9997 |
| | **75 (3)[e]** | 51.63 ± 1.83 | 0.0041 | 53.26 ± 2.59 | 0.0008 | 1.0000 | 52.30 ± 2.05 | 0.0002 | 51.66 ± 4.80 | 0.0117 | 1.0000 | 1.0000 | 1.0000 |
| | **77 (5)[e]** | 53.73 ± 1.95 | 0.0017 | 54.22 ± 1.23 | 0.0003 | 1.0000 | 53.75 ± 2.52 | <.0001 | 48.35 ± 4.23 | 0.0300 | 0.9628 | 1.0000 | 0.9477 |

[a]Mean percentage of MLN cells ± SEM.
[b]p-value calculated by Dunnett's post-hoc test (day relative to primary infection compared to day 0 within sex/developmental exposure group).
[c]p-value calculated by Tukey HSD post-hoc test (PFAS vs. CTRL within sex on each day relative to primary infection).
[d]p-value calculated by Tukey HSD post-hoc test (male vs. female within developmental exposure group on each day relative to primary infection).
[e]Number in parenthesis () indicates day relative to secondary infection.

**Table S6.** Percentage of CD4+ T cell subsets in the MLN.

| Cell type | Day relative to primary infection | Male CTRL[a] | p-value[b] | Male PFAS[a] | p-value[b] | p-value[c] | Female CTRL[a] | p-value[b] | Female PFAS[a] | p-value[b] | p-value[c] | p-value CTRL male vs. CTRL female[d] | p-value PFAS male vs. PFAS female[d] |
|---|---|---|---|---|---|---|---|---|---|---|---|---|---|
| Th1 | 0 | 2.16 ± 0.53 | 1.0000 | 2.55 ± 0.34 | 1.0000 | 1.0000 | 2.08 ± 0.20 | 1.0000 | 2.55 ± 0.34 | 1.0000 | 1.0000 | 1.0000 | 0.9997 |
| | 9 | 5.86 ± 1.13 | 0.0378 | 5.43 ± 0.12 | 0.0922 | 1.0000 | 6.60 ± 1.10 | 0.0068 | 5.43 ± 0.12 | 0.0922 | 0.9903 | 0.9999 | 0.9981 |
| | 70 | 3.77 ± 0.40 | 0.4839 | 3.99 ± 0.94 | 0.7071 | 0.9978 | 3.26 ± 0.51 | 0.8164 | 3.99 ± 0.94 | 0.7071 | 1.0000 | 1.0000 | 0.9972 |
| | 75 (3)[e] | 7.44 ± 0.77 | 0.0006 | 10.74 ± 1.06 | <.0001 | 1.0000 | 9.84 ± 1.36 | <.0001 | 10.74 ± 1.06 | <.0001 | 0.9991 | 0.4700 | 0.0930 |
| | 77 (5)[e] | 4.36 ± 0.88 | 0.2175 | 5.11 ± 1.14 | 0.1480 | 0.9997 | 4.68 ± 0.33 | 0.1630 | 5.11 ± 1.14 | 0.1480 | 1.0000 | 1.0000 | 0.9509 |
| Th2 | 0 | 0.17 ± 0.02 | 1.0000 | 0.32 ± 0.14 | 1.0000 | 0.9975 | 0.13 ± 0.02 | 1.0000 | 0.32 ± 0.14 | 1.0000 | 0.9970 | 1.0000 | 1.0000 |
| | 9 | 0.25 ± 0.07 | 0.9550 | 0.29 ± 0.03 | 0.9999 | 1.0000 | 0.29 ± 0.04 | 0.6781 | 0.29 ± 0.03 | 0.9999 | 1.0000 | 1.0000 | 1.0000 |
| | 70 | 0.61 ± 0.09 | 0.0199 | 0.54 ± 0.07 | 0.8719 | 0.7242 | 0.51 ± 0.07 | 0.1021 | 0.54 ± 0.07 | 0.8719 | 1.0000 | 0.9998 | 0.9984 |
| | 75 (3)[e] | 0.54 ± 0.12 | 0.0385 | 0.95 ± 0.15 | 0.0964 | 0.9726 | 0.72 ± 0.15 | 0.0031 | 0.95 ± 0.16 | 0.0964 | 0.9772 | 0.9351 | 0.9590 |
| | 77 (5)[e] | 0.95 ± 0.06 | <.0001 | 1.70 ± 0.28 | 0.0002 | 0.9132 | 1.75 ± 0.15 | <.0001 | 1.70 ± 0.28 | 0.0002 | 1.0000 | <.0001 | 0.0977 |
| Th17 | 0 | 0.60 ± 0.10 | 1.0000 | 3.36 ± 1.54 | 1.0000 | 1.0000 | 0.84 ± 0.11 | 1.0000 | 3.36 ± 1.54 | 1.0000 | 0.0186 | 0.9997 | 0.1187 |
| | 9 | 0.93 ± 0.32 | 0.8951 | 0.79 ± 0.07 | 0.0384 | 1.0000 | 0.74 ± 0.09 | 0.9619 | 0.79 ± 0.07 | 0.0384 | 1.0000 | 0.9999 | 1.0000 |
| | 70 | 1.77 ± 0.09 | 0.0556 | 1.11 ± 0.05 | 0.1568 | 1.0000 | 1.13 ± 0.26 | 0.5293 | 1.11 ± 0.05 | 0.1568 | 1.0000 | 0.7675 | 0.9972 |
| | 75 (3)[e] | 2.09 ± 0.41 | 0.0079 | 1.83 ± 0.25 | 0.3357 | 1.0000 | 1.49 ± 0.20 | 0.0161 | 1.83 ± 0.25 | 0.3357 | 0.9999 | 0.6342 | 1.0000 |
| | 77 (5)[e] | 1.66 ± 0.18 | 0.0777 | 2.36 ± 0.56 | 0.6449 | 0.9530 | 1.63 ± 0.06 | 0.0033 | 2.36 ± 0.56 | 0.6449 | 0.9596 | 1.0000 | 1.0000 |
| Treg | 0 | 10.54 ± 1.05 | 1.0000 | 9.06 ± 0.82 | 1.0000 | 1.0000 | 10.10 ± 0.39 | 1.0000 | 9.06 ± 0.82 | 1.0000 | 0.9949 | 1.0000 | 1.0000 |
| | 9 | 10.98 ± 2.54 | 0.9989 | 10.68 ± 0.35 | 0.3737 | 1.0000 | 9.12 ± 0.35 | 0.7407 | 10.68 ± 0.35 | 0.3737 | 0.8375 | 0.9861 | 1.0000 |
| | 70 | 14.75 ± 0.91 | 0.1791 | 10.69 ± 1.01 | 0.5047 | 0.9970 | 12.10 ± 1.87 | 0.2676 | 10.69 ± 1.01 | 0.5047 | 0.9829 | 0.8798 | 0.8387 |
| | 75 (3)[e] | 12.73 ± 1.44 | 0.6408 | 11.31 ± 0.93 | 0.1618 | 1.0000 | 10.77 ± 0.72 | 0.9111 | 11.31 ± 0.93 | 0.1618 | 0.9999 | 0.9400 | 0.9555 |
| | 77 (5)[e] | 9.00 ± 1.00 | 0.8547 | 7.31 ± 0.74 | 0.3085 | 1.0000 | 8.58 ± 0.14 | 0.4051 | 7.31 ± 0.74 | 0.3085 | 0.9465 | 1.0000 | 0.9848 |
| Tfh | 0 | 1.44 ± 0.47 | 1.0000 | 3.49 ± 1.47 | 1.0000 | 0.8558 | 2.55 ± 0.70 | 1.0000 | 4.64 ± 0.77 | 1.0000 | 0.8703 | 0.9947 | 0.9986 |
| | 9 | 7.83 ± 1.19 | <.0001 | 6.70 ± 0.67 | 0.1222 | 0.9938 | 6.97 ± 1.56 | 0.0084 | 5.52 ± 1.07 | 0.8762 | 0.9823 | 0.9990 | 0.9927 |
| | 70 | 4.35 ± 0.66 | 0.0099 | 7.39 ± 2.22 | 0.0778 | 0.2698 | 2.77 ± 0.79 | 0.9996 | 5.33 ± 1.58 | 0.9667 | 0.8152 | 0.9260 | 0.9542 |
| | 75 (3)[e] | 3.23 ± 0.35 | 0.1216 | 3.15 ± 0.38 | 0.9979 | 1.0000 | 3.67 ± 0.34 | 0.7943 | 2.63 ± 0.58 | 0.3501 | 0.9975 | 1.0000 | 1.0000 |
| | 77 (5)[e] | 3.33 ± 0.35 | 0.1075 | 2.57 ± 0.37 | 0.9153 | 0.9991 | 4.24 ± 0.45 | 0.5012 | 2.56 ± 0.47 | 0.2925 | 0.9128 | 0.9949 | 1.0000 |

Th1: $Tbet^+CD4^+$; Th2: $GATA3^+CD4^+$; Th17: $ROR\gamma t^+CD4^+$; Treg: $FoxP3^+CD25^+CD4^+$; Tfh: $PD\text{-}1^{hi}CXCR5^+CD44^{hi}CD4^+$

[a]Mean percentage of CD4+ T cells ± SEM.

[b]p-value calculated by Dunnett's post-hoc test (day relative to primary infection compared to day 0 within sex/developmental exposure group).

[c]p-value calculated by Tukey HSD post-hoc test (PFAS vs. CTRL within sex on each day relative to primary infection).

[d]p-value calculated by Tukey HSD post-hoc test (male vs. female with developmental exposure group on each day relative to primary infection).

[e]Number in parenthesis () indicates day relative to secondary infection.

**Table S7.** Number of cells in CD4+ T cell subsets in the MLN.

| Cell type | Day relative to primary infection | Male CTRL[a] | p-value[b] | Male PFAS[a] | p-value[b] | p-value[c] | Female CTRL[a] | p-value[b] | Female PFAS[a] | p-value[b] | p-value[c] | p-value CTRL male vs. CTRL female[d] | p-value PFAS male vs. PFAS female[d] |
|---|---|---|---|---|---|---|---|---|---|---|---|---|---|
| Th1 | 0 | 9491 ± 3605 | 1.0000 | 4272 ± 1361 | 1.0000 | 0.9999 | 7994 ± 1583 | 1.0000 | 14342 ± 5216 | 1.0000 | 1.0000 | 1.0000 | 0.9997 |
| | 9 | 45051 ± 11018 | 0.0075 | 37227 ± 2894 | <.0001 | 0.9919 | 49308 ± 13130 | 0.0292 | 40249 ± 5033 | 0.4329 | 0.9998 | 1.0000 | 1.0000 |
| | 70 | 10693 ± 3688 | 0.9997 | 7035 ± 1728 | 0.9705 | 1.0000 | 13626 ± 4851 | 0.9889 | 26332 ± 12273 | 0.9435 | 0.9997 | 1.0000 | 0.9672 |
| | 75 (3)[e] | 29364 ± 5868 | 0.1069 | 22697 ± 5633 | 0.0233 | 0.9937 | 61821 ± 10965 | 0.0040 | 69181 ± 22657 | 0.0313 | 1.0000 | 0.0915 | 0.0349 |
| | 77 (5)[e] | 20521 ± 5446 | 0.5462 | 13296 ± 3770 | 0.3524 | 0.9868 | 39688 ± 10455 | 0.1162 | 34078 ± 10185 | 0.6503 | 1.0000 | 0.7545 | 0.7800 |
| Th2 | 0 | 682 ± 227 | 1.0000 | 720 ± 215 | 1.0000 | 1.0000 | 432 ± 16 | 1.0000 | 1191 ± 265 | 1.0000 | 1.0000 | 1.0000 | 1.0000 |
| | 9 | 1853 ± 552 | 0.3818 | 1637 ± 244 | 0.3893 | 1.0000 | 1868 ± 347 | 0.8026 | 2146 ± 364 | 0.9784 | 1.0000 | 1.0000 | 1.0000 |
| | 70 | 1582 ± 565 | 0.5224 | 885 ± 269 | 0.9965 | 0.9876 | 2029 ± 557 | 0.8007 | 3375 ± 1436 | 0.8252 | 0.9999 | 1.0000 | 0.9571 |
| | 75 (3)[e] | 1708 ± 268 | 0.3724 | 2280 ± 434 | 0.0804 | 0.9926 | 4260 ± 892 | 0.1013 | 6357 ± 2160 | 0.1272 | 0.9825 | 0.4304 | 0.3080 |
| | 77 (5)[e] | 4124 ± 553 | 0.0002 | 3582 ± 539 | 0.0008 | 0.9940 | 13630 ± 2148 | <.0001 | 10253 ± 2079 | 0.0030 | 0.7005 | <.0001 | 0.0026 |
| Th17 | 0 | 2837 ± 1188 | 1.0000 | 1987 ± 208 | 1.0000 | 1.0000 | 3157 ± 570 | 1.0000 | 12417 ± 3837 | 1.0000 | 0.0650 | 1.0000 | 0.0302 |
| | 9 | 6837 ± 2509 | 0.1755 | 4631 ± 546 | 0.1592 | 0.9319 | 4643 ± 551 | 0.8758 | 5794 ± 943 | 0.1671 | 1.0000 | 0.9846 | 0.9999 |
| | 70 | 4688 ± 1369 | 0.6916 | 4651 ± 667 | 0.2008 | 1.0000 | 4770 ± 1975 | 0.8830 | 6249 ± 2053 | 0.3371 | 1.0000 | 1.0000 | 0.9999 |
| | 75 (3)[e] | 6639 ± 617 | 0.1238 | 6156 ± 1405 | 0.0201 | 1.0000 | 8806 ± 818 | 0.0384 | 10884 ± 2459 | 0.9720 | 0.9981 | 0.9549 | 0.6374 |
| | 77 (5)[e] | 7053 ± 900 | 0.0876 | 6348 ± 642 | 0.0118 | 1.0000 | 12929 ± 2387 | 0.0004 | 13015 ± 2185 | 0.9991 | 1.0000 | 0.0469 | 0.1115 |
| Treg | 0 | 47381 ± 18047 | 1.0000 | 24137 ± 4006 | 1.0000 | 0.9499 | 42451 ± 11888 | 1.0000 | 57949 ± 21755 | 1.0000 | 0.9997 | 1.0000 | 0.8932 |
| | 9 | 83859 ± 23819 | 0.1934 | 64944 ± 8784 | 0.0121 | 0.9602 | 65234 ± 15030 | 0.6309 | 78410 ± 8556 | 0.8156 | 0.9997 | 0.9940 | 0.9985 |
| | 70 | 40608 ± 11859 | 0.9818 | 33193 ± 5261 | 0.8743 | 1.0000 | 47440 ± 12511 | 0.9982 | 63993 ± 23661 | 0.9985 | 0.9998 | 1.0000 | 0.9361 |
| | 75 (3)[e] | 44250 ± 5175 | 0.9988 | 44759 ± 10363 | 0.3032 | 1.0000 | 72452 ± 14796 | 0.4041 | 77826 ± 24509 | 0.8463 | 1.0000 | 0.7859 | 0.7644 |
| | 77 (5)[e] | 38999 ± 6062 | 0.9577 | 28862 ± 5657 | 0.9803 | 0.9986 | 70355 ± 15308 | 0.4656 | 52922 ± 13017 | 0.9986 | 0.9975 | 0.7057 | 0.9231 |
| Tfh | 0 | 345 ± 240 | 1.0000 | 343 ± 195 | 1.0000 | 1.0000 | 1095 ± 497 | 1.0000 | 2445 ± 858 | 1.0000 | 1.0000 | 1.0000 | 0.9953 |
| | 9 | 10975 ± 3513 | 0.0003 | 7806 ± 1315 | <.0001 | 0.7092 | 13633 ± 4217 | 0.0051 | 7363 ± 2476 | 0.2541 | 0.5181 | 0.9957 | 1.0000 |
| | 70 | 2254 ± 845 | 0.7349 | 2429 ± 626 | 0.3138 | 1.0000 | 1785 ± 765 | 0.9992 | 5940 ± 2869 | 0.6597 | 0.9853 | 1.0000 | 0.9194 |
| | 75 (3)[e] | 3327 ± 1028 | 0.3536 | 2277 ± 265 | 0.3320 | 0.9995 | 5977 ± 1155 | 0.4339 | 4058 ± 1427 | 0.9415 | 0.9998 | 0.9849 | 0.9969 |
| | 77 (5)[e] | 3494 ± 666 | 0.3288 | 1991 ± 509 | 0.4309 | 0.9901 | 10076 ± 2058 | 0.0520 | 4421 ± 1532 | 0.8768 | 0.6554 | 0.2519 | 0.9516 |

Th1: Tbet$^+$CD4$^+$; Th2: GATA3$^+$CD4$^+$; Th17: ROR$\gamma$t$^+$CD4$^+$; Treg: FoxP3$^+$CD25$^+$CD4$^+$; Tfh: PD-1$^{hi}$CXCR5$^+$CD44$^{hi}$CD4$^+$

[a]Mean number of cells ± SEM.

[b]p-value calculated by Dunnett's post-hoc test (day relative to primary infection compared to day 0 within sex/developmental exposure group).

[c]p-value calculated by Tukey HSD post-hoc test (PFAS vs. CTRL within sex on each day relative to primary infection).

[d]p-value calculated by Tukey HSD post-hoc test (male vs. female with developmental exposure group on each day relative to primary infection).

[e]Number in parenthesis () indicates day relative to secondary infection.

**Table S8.** Percentage and number of germinal center (GC) B cells and plasma cells (PC) in the MLN.

| Cell type | Day relative to primary infection | Male CTRL[a] | p-value[b] | Male PFAS[a] | p-value[b] | p-value[c] | Female CTRL[a] | p-value[b] | Female PFAS[a] | p-value[b] | p-value[c] | p-value CTRL male vs. CTRL female[d] | p-value PFAS male vs. PFAS female[d] |
|---|---|---|---|---|---|---|---|---|---|---|---|---|---|
| % GC B cells | 0 | 0.004 ± 0.0003 | 1.0000 | 0.01 ± 0.003 | 1.0000 | 1.0000 | 0.01 ± 0.003 | 1.0000 | 0.008 ± 0.003 | 1.0000 | 1.0000 | 1.0000 | 1.0000 |
| | 14 | 0.62 ± 0.16 | 0.0149 | 0.63 ± 0.13 | 0.0020 | 1.0000 | 0.40 ± 0.03 | 0.0371 | 0.34 ± 0.06 | 0.0129 | 1.0000 | 0.9098 | 0.3105 |
| | 70 | 0.62 ± 0.19 | 0.0157 | 0.41 ± 0.12 | 0.0767 | 0.9756 | 0.51 ± 0.18 | 0.0056 | 0.57 ± 0.16 | 0.0006 | 1.0000 | 0.9995 | 0.9897 |
| | 75 (3)[e] | 0.12 ± 0.04 | 0.9064 | 0.20 ± 0.09 | 0.5508 | 1.0000 | 0.15 ± 0.08 | 0.7177 | 0.08 ± 0.03 | 0.8769 | 1.0000 | 1.0000 | 0.9954 |
| | 77 (5)[e] | 0.51 ± 0.11 | 0.0456 | 0.40 ± 0.09 | 0.0538 | 0.9995 | 0.66 ± 0.06 | 0.0004 | 0.41 ± 0.09 | 0.0027 | 0.5142 | 0.9912 | 1.0000 |
| # GC B cells | 0 | 70 ± 24 | 1.0000 | 173 ± 42 | 1.0000 | 1.0000 | 153 ± 32 | 1.0000 | 150 ± 71 | 1.0000 | 1.0000 | 1.0000 | 1.0000 |
| | 14 | 30332 ± 13112 | 0.0260 | 25014 ± 7797 | 0.0058 | 0.9996 | 20071 ± 4745 | 0.0225 | 12493 ± 4305 | 0.1591 | 0.9611 | 0.9566 | 0.5301 |
| | 70 | 13902 ± 5784 | 0.4659 | 7430 ± 2986 | 0.7173 | 1.0000 | 13401 ± 6920 | 0.1739 | 20707 ± 11769 | 0.0337 | 0.9922 | 1.0000 | 0.7927 |
| | 75 (3)[e] | 3357 ± 937 | 0.9901 | 5306 ± 2350 | 0.8676 | 0.9997 | 6697 ± 2119 | 0.7184 | 4694 ± 1732 | 0.8796 | 1.0000 | 1.0000 | 1.0000 |
| | 77 (5)[e] | 14936 ± 3455 | 0.3832 | 9873 ± 2670 | 0.4239 | 1.0000 | 35935 ± 4877 | <.0001 | 18097 ± 4141 | 0.0244 | 0.1302 | 0.2070 | 0.9240 |
| % PC | 0 | 0.07 ± 0.01 | 1.0000 | 0.09 ± 0.03 | 1.0000 | 1.0000 | 0.18 ± 0.04 | 1.0000 | 0.20 ± 0.03 | 1.0000 | 0.9999 | 0.7617 | 0.8667 |
| | 14 | 0.19 ± 0.05 | 0.2347 | 0.27 ± 0.06 | 0.1535 | 0.9691 | 0.23 ± 0.02 | 0.6227 | 0.23 ± 0.04 | 0.9365 | 1.0000 | 0.9989 | 0.9998 |
| | 70 | 0.33 ± 0.04 | 0.0022 | 0.42 ± 0.13 | 0.0073 | 0.9578 | 0.26 ± 0.05 | 0.1841 | 0.27 ± 0.06 | 0.5541 | 1.0000 | 0.9560 | 0.6858 |
| | 75 (3)[e] | 0.19 ± 0.05 | 0.1843 | 0.18 ± 0.04 | 0.7484 | 1.0000 | 0.12 ± 0.02 | 0.4793 | 0.14 ± 0.02 | 0.4860 | 0.9999 | 0.8734 | 1.0000 |
| | 77 (5)[e] | 0.17 ± 0.03 | 0.3365 | 0.16 ± 0.02 | 0.8364 | 1.0000 | 0.12 ± 0.01 | 0.5058 | 0.13 ± 0.01 | 0.2977 | 1.0000 | 0.9893 | 1.0000 |
| # PC | 0 | 1466 ± 692 | 1.0000 | 2050 ± 1079 | 1.0000 | 1.0000 | 2612 ± 686 | 1.0000 | 4349 ± 1505 | 1.0000 | 0.9993 | 1.0000 | 0.9948 |
| | 14 | 8632 ± 3947 | 0.0810 | 8793 ± 2318 | 0.0524 | 1.0000 | 11215 ± 2946 | 0.0087 | 8097 ± 1838 | 0.2435 | 0.9221 | 0.9913 | 1.0000 |
| | 70 | 5685 ± 1146 | 0.4265 | 7179 ± 2730 | 0.2320 | 0.9999 | 5616 ± 1260 | 0.5786 | 8054 ± 1823 | 0.4032 | 0.9960 | 1.0000 | 1.0000 |
| | 75 (3)[e] | 5522 ± 708 | 0.4346 | 4387 ± 1197 | 0.7808 | 1.0000 | 6848 ± 1172 | 0.2945 | 7846 ± 1516 | 0.3313 | 1.0000 | 0.9999 | 0.8963 |
| | 77 (5)[e] | 4958 ± 879 | 0.5558 | 3946 ± 965 | 0.8612 | 1.0000 | 6940 ± 1496 | 0.2780 | 5261 ± 830 | 0.9766 | 0.9989 | 0.9984 | 0.9998 |

GC B cells: $CD3^{-}B220^{+}CD95^{+}GL7^{+}$; PC: $CD3^{-}B220^{+/-}CD138^{+}$

[a]Mean percentage of MLN cells/mean number of cells ± SEM.

[b]p-value calculated by Dunnett's post-hoc test (day relative to primary infection compared to day 0 within sex/developmental exposure group).

[c]p-value calculated by Tukey HSD post-hoc test (PFAS vs. CTRL within sex on each day relative to primary infection).

[d]p-value calculated by Tukey HSD post-hoc test (male vs. female with developmental exposure group on each day relative to primary infection).

[e]Number in parenthesis () indicates day relative to secondary infection.

**Table S9.** Area under curve (AUC) values and statistical comparisons for IAV-specific IgM and IgG.

| Antibody | Day relative to primary infection | Male CTRL[a] | p-value[b] | Male PFAS[a] | p-value[b] | p-value[c] | Female CTRL[a] | p-value[b] | Female PFAS[a] | p-value[b] | p-value[c] | p-value CTRL male vs. CTRL female[d] | p-value PFAS male vs. PFAS female[d] |
|---|---|---|---|---|---|---|---|---|---|---|---|---|---|
| IgM | **0** | 0.37 ± 0.08 | 1.0000 | 0.36 ± 0.04 | 1.0000 | >0.9999 | 0.30 ± 0.05 | 1.0000 | 0.32 ± 0.04 | 1.0000 | >0.9999 | >0.9999 | >0.9999 |
| | **14** | 2.34 ± 0.55 | <0.0001 | 2.19 ± 0.62 | <0.0001 | >0.9999 | 2.93 ± 0.33 | <0.0001 | 3.93 ± 0.47 | <0.0001 | 0.1867 | 0.8477 | 0.0008 |
| | **70** | 0.62 ± 0.10 | 0.9168 | 0.53 ± 0.05 | 0.9784 | >0.9999 | 2.23 ± 0.27 | <0.0001 | 1.85 ± 0.17 | 0.0030 | 0.9950 | 0.0003 | 0.0347 |
| | **75 (3)[e]** | 0.80 ± 0.11 | 0.6660 | 1.00 ± 0.20 | 0.2634 | >0.9999 | 2.19 ± 0.30 | <0.0001 | 2.44 ± 0.27 | <0.0001 | 0.9997 | 0.0083 | 0.0022 |
| | **77 (5)[e]** | 1.27 ± 0.16 | 0.1231 | 0.76 ± 0.11 | 0.6515 | 0.9112 | 1.15 ± 0.11 | 0.0972 | 0.98 ± 0.08 | 0.4856 | >0.9999 | >0.9999 | >0.9999 |
| IgG | **0** | 0.07 ± 0.02 | 1.0000 | 0.09 ± 0.02 | 1.0000 | >0.9999 | 0.10 ± 0.10 | 1.0000 | 0.03 ± 0.01 | 1.0000 | >0.9999 | >0.9999 | >0.9999 |
| | **14** | 6.80 ± 0.28 | <0.0001 | 6.40 ± 0.29 | <0.0001 | 0.9787 | 7.23 ± 0.19 | <0.0001 | 7.80 ± 0.24 | <0.0001 | 0.7709 | 0.9556 | 0.0028 |
| | **70** | 5.36 ± 0.19 | <0.0001 | 4.69 ± 0.11 | <0.0001 | 0.4767 | 7.08 ± 0.29 | <0.0001 | 5.94 ± 0.22 | <0.0001 | 0.0534 | <0.0001 | 0.0137 |
| | **75 (3)[e]** | 5.68 ± 0.25 | <0.0001 | 6.22 ± 0.32 | <0.0001 | 0.7592 | 7.42 ± 0.28 | <0.0001 | 7.59 ± 0.32 | <0.0001 | >0.9999 | <0.0001 | 0.0005 |
| | **77 (5)[e]** | 6.01 ± 0.25 | <0.0001 | 4.40 ± 0.09 | <0.0001 | <0.0001 | 7.09 ± 0.21 | <0.0001 | 5.63 ± 0.29 | <0.0001 | 0.0015 | 0.0409 | 0.0029 |

[a]Mean area under curve ± SEM.
[b]p-value calculated by Dunnett's post-hoc test (day relative to primary infection vs. day 0 within sex/developmental exposure group).
[c]p-value calculated by Tukey HSD post-hoc test (PFAS vs. CTRL within sex on each day relative to primary infection).
[d]p-value calculated by Tukey HSD post-hoc test (male vs. female within developmental exposure group on each day relative to primary infection).
[e]Number in parenthesis () indicates day relative to secondary infection.

**Table S10**. Avidity ELISA area under curve analysis statistical comparisons.

| Sex | Group 1 | AUC | Group 2 | AUC | p-value[a] |
|---|---|---|---|---|---|
| **Male** | CTRL -GuHCl | 6.012 | CTRL +GuHCl | 4.637 | <0.0001 |
| | CTRL -GuHCl | 6.012 | PFAS -GuHCl | 4.395 | <0.0001 |
| | CTRL +GuHCl | 4.637 | PFAS +GuHCl | 3.315 | <0.0001 |
| | PFAS -GuHCl | 4.395 | PFAS +GuHCl | 3.315 | 0.0003 |
| **Female** | CTRL -GuHCl | 7.088 | CTRL +GuHCl | 6.682 | 0.6601 |
| | CTRL -GuHCl | 7.088 | PFAS -GuHCl | 5.628 | 0.0004 |
| | CTRL +GuHCl | 6.682 | PFAS +GuHCl | 4.706 | <0.0001 |
| | PFAS -GuHCl | 5.628 | PFAS +GuHCl | 4.706 | 0.0499 |

[a]p-value calculated by Tukey HSD post-hoc test.
